# Innovations in spinal cord cell type heterogeneity across vertebrate evolution

**DOI:** 10.1101/2025.10.09.680955

**Authors:** Yuri Ignatyev, Stavros Papadopoulos, Mateja Soretić, Jake Yeung, Tzi-Yang Lin, Elly M Tanaka, Leonid Peshkin, Ariel J Levine, Mariano I Gabitto, Lora B Sweeney

## Abstract

Vertebrates display remarkable diversity of sensorimotor behaviors, each adapted to distinct ecological and survival demands. This diversity raises fundamental questions about the evolutionary origin of motor control: do conserved spinal circuits underlie these behaviors, and how have they diverged across species. Recent studies detail spinal cell-type architecture in mammals but comparable, high-resolution atlases of the non-mammalian spinal cord are lacking. Here, we compare spinal cord cell types between fish, frogs, mice and humans, spanning ∼450 million years of evolution. Across species, we define highly conserved programs of cell type specification that segregate spinal neurons into nearly identical cardinal classes during development. This contrasts with adult stages, when spinal cell-type composition selectively diverges for excitatory neuron subpopulations. Using spatial transcriptomics, we localize this species divergence to the superficial, dorsal spinal cord, where variant neuropeptide expression defines mammalian-specific cell types. The most dorsal spinal cord thus emerges as a recently evolved hub for sensory integration in mammals, a neospinal cord analogous to the neocortex.

## Main Text

During evolution, vertebrates transitioned from an aquatic to terrestrial environment and correspondingly evolved four limbs.^1^ Terrestrial environments had stronger gravitational forces, and more variant topography and temperatures, and thus land-based movement corresponded with an increase in fine-tuned proprioception and precision of nociceptive, tactile, and noxious sensory processing.^2,3^ In addition, with limbs, greater precision, control and adaptability of movement became possible, exemplified by dexterous, voluntary movements such as grasping.^4–6^ Such variations in movement execution, sensory processing, and the integration of ascending and descending commands to and from the brain are executed by spinal circuits.^7^ However, despite decades of research on spinal circuit composition, we almost completely lack an understanding of how these dramatic changes in sensorimotor processing were reflected in the evolution of vertebrate spinal circuits. This knowledge gap hinders our ability to link spinal cord architecture, and specific neuron types, with the diversity of organismal movement and sensory processing, and leaves open the question of what mechanism drove the evolutionary diversification of vertebrate motor behavior.

The intricate and well-described spinal cell types of the mammalian spinal cord provide a starting point to define shared and divergent circuit properties across evolutionary time. During development, gene regulatory programs in the mouse, primate and human spinal cord^8–16^ segregate spinal neurons into eleven cardinal classes, each characterized by unique transcription factor expression, neurotransmitter profile, projection pattern, and function.^1^ Within each of these cardinal classes, the level of cell-type heterogeneity is immense and in most cases, the function of this heterogeneity is unknown^1,17–19^: inhibitory V1 interneurons, for example, can each contain as many as 50 molecularly-distinct subtypes.^20–22^ While individual examples exist of molecularly conserved cardinal classes in zebrafish,^23–27^ the extent to which this cardinal architecture in its entirety extends beyond mammals — and how it may vary with species-specific movement and sensation — is unknown. Still open too is the question of whether fine-scale subtype heterogeneity is a uniquely mammalian trait or a more ancient vertebrate feature.

In addition, while this cardinal architecture defines the developing mammalian spinal cord, in adults, the defining markers of cardinal identity are downregulated. The mature, adult spinal cord in contrast is organized into a ten-layer laminar structure along the dorsal-ventral axis, with individual neuron types displaying highly specific distribution patterns and distinct gene expression signatures from those in development.^12,28,29^ Mature ventral spinal neurons striate into excitatory and inhibitory subpopulations, which initiate or shape the motor reflex pattern.^14,15,19^ Dorsal neuron types, which include many molecularly heterogeneous subpopulations, are distributed across laminae I-V in accordance with the neuropeptides they express and the sensory function they regulate^30^. Despite this detailed knowledge of cell type identity and distribution in mammals however, the conservation of this adult architecture in other vertebrates is almost completely uncharacterized — leaving still open the basic question of how this architecture evolved, and whether it relates to the specialized sensorimotor processing capabilities of mammals.

Here, we take a cross-species comparative approach to study vertebrate spinal cord evolution. We capitalize on the evolutionary position of amphibians, at the transition point between swimming and limbed motor behavior, to evaluate conservation and divergence of spinal cord cell types across ∼450 million years of evolution, spanning teleost fish, amphibians and mammals (**Fig. 1A**). Between these distantly related species, we find deep conservation of the cardinal class architecture during development, indicating high convergence of early spinal cord specification. This sharply contrasts with adult stages, in which we discover mammalian-specific innovations of mature dorsal excitatory neuron types, absent from both fish and frogs. The most superficial dorsal spinal cord thus emerges from our analysis as a recently evolved hub for sensory integration that is unique to mammals, a neospinal cord analogous to the neocortex.

**Fig. 1.**
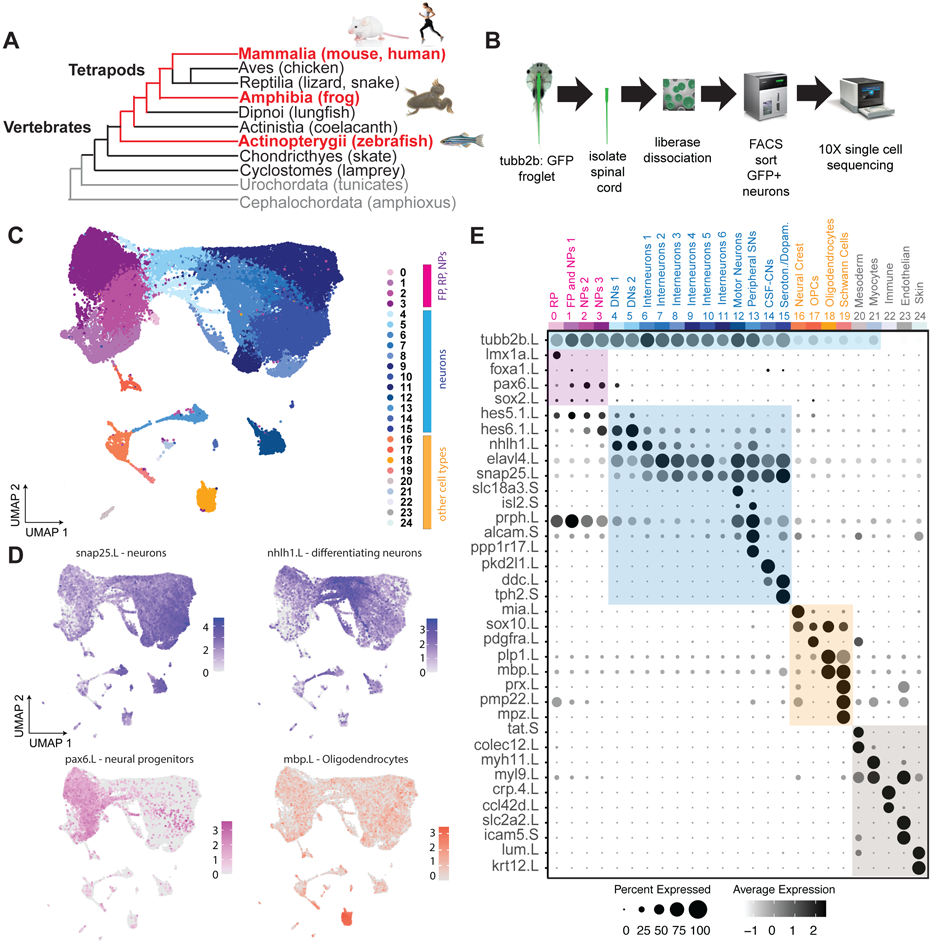
Transcriptomic cell-type atlas of the developing frog spinal cord. **A.** Schematic phylogeny of *Chordates* highlighting in red the organisms compared in this study: the four-limbed amphibian *Xenopus laevis* (frog), actinopterygii (zebrafish), and mammals (mouse and human). **B.** Experimental workflow including isolation, enrichment, and sequencing of frog spinal neurons. Transgenic froglets expressing GFP under the *tubb2b* promoter were dissected, and their spinal neurons isolated for analysis. **C.** UMAP representation of ∼39,000 frog spinal transcriptomes color coded by cell type after sequencing and data processing. Numbers denote clusters identities defined by unsupervised clustering. **D**. Feature plots demonstrating the expression level in each cell of pan-neuronal marker, *snap25*; differentiating neuron marker, *nhlh1l;* neural progenitor marker, *pax6*; and oligodendrocyte marker, *mbp*. **E**. Dot plot showing the expression of cell type–specific markers across populations defined in **C**. Dot size indicates the percentage of cells expressing each marker per cluster, and color denotes the average scaled expression level. Abbreviations: RF, roof plate; FP, floor plate; NP, neural progenitors; OPC, oligodendrocyte progenitor cells; DN, differentiating neurons; SN, sensory neurons; CSF-CNs, cerebrospinal fluid–contacting neurons.

### A cell-type atlas of the developing frog spinal cord

The basic cell-type architecture for movement is established during mammalian spinal cord development,^31^ motivating us to characterize the developing *Xenopus laevis* frog spinal cord. To evaluate cell type diversity, we generated a single-cell atlas at the peak of frog limb circuit development, NF^32^ stage 54 (NF54), when the limb and all limb-level motor neurons and interneurons have been generated.^33^ Spinal neurons were obtained by FACS sorting GFP-positive cells from isolated spinal cords of pan-neuronal *tubb2b-GFP*^34^ transgenic animals, and single-cell RNA sequencing was performed (scRNA-seq, 10X Genomics; **Fig. 1B**). Reads were mapped to *X. laevis* reference transcriptome (see *Methods*). After quality filtering (**Fig. S1**), 39,338 cells were obtained with an average of ∼2600 genes detected per cell, aligning with previous scRNA-sequencing studies of the *Xenopus* tail.^35^ Unsupervised clustering analysis identified major spinal cord and associated cell types including neural progenitors, neurons, neural crest and glial cells (**Fig. 1C**). Cell identities of these clusters were determined based on marker gene expression: *snap25/elavl4* for neurons, *nhlh1/hes6 for* differentiating, immature neurons*, pax6/sox2* for neural progenitors, and *mbp/pdgfra* for oligodendrocytes (**Fig. 1D-E**).

Mature (*snap25-*expressing) and differentiating (*nhlh*1-expressing) neurons were the most prominent groups, together constituting ∼60% of our data. Neural progenitors (*sox2/pax6*-expressing) made up ∼30%. A variety of other cell types, including roof and floor plate, neural crest, glia, muscle, and skin cells (**Fig. 1E**), accounted for the remaining ∼10% of total cells — confirming the success of our FACS-based strategy to enrich for neurons. Clusters of mature neurons included ∼22,000 single-cell profiles composed of heterogeneous interneurons, motor neurons, and small fractions of peripheral sensory neurons, cerebrospinal fluid-contacting neurons, and serotonergic/dopaminergic neurons (**Fig. 1E**). Subsequent analysis focused on motor and interneurons, due to their central role in the generation and regulation of vertebrate movement patterns.

### Conserved developmental spinal cell-type architecture across evolutionarily distant tetrapods

The hallmark cell-type architecture of the developing mammalian spinal cord includes cardinal classes of inhibitory and excitatory interneurons.^36^ This cardinal class structure delineates anatomically and functionally distinct interneuron types that together orchestrate motor and sensory processing.^19^ To evaluate whether cardinal cell type architecture is conserved in amphibians, we compared our NF54 *Xenopus* frog spinal cord atlas with an analogous neuronal atlas from developing mice.^13^ Comparable developmental stages between mouse and frog were identified via alignment of landmarks of motor, spinal cord and limb development across species^37^ and further corroborated by gene expression. This enabled a detailed comparison of spinal neurons and their transcriptional determinants between species. To assess conservation of cardinal class architecture, we applied high-resolution Louvain clustering separately to frog and mouse (**Fig. S2**) neuronal data. We performed differential expression analysis to identify cluster-specific genes and annotated each cluster based on the expression of established markers.^13^ All major cardinal classes were identified in the frog spinal cord, with an identical core set of transcription factors segregating each inhibitory and excitatory cardinal class in both frogs and mice (**Fig. 2A-B**), and each class also comprising similar cellular proportions across species (**Fig. 2C).**

**Fig. 2.**
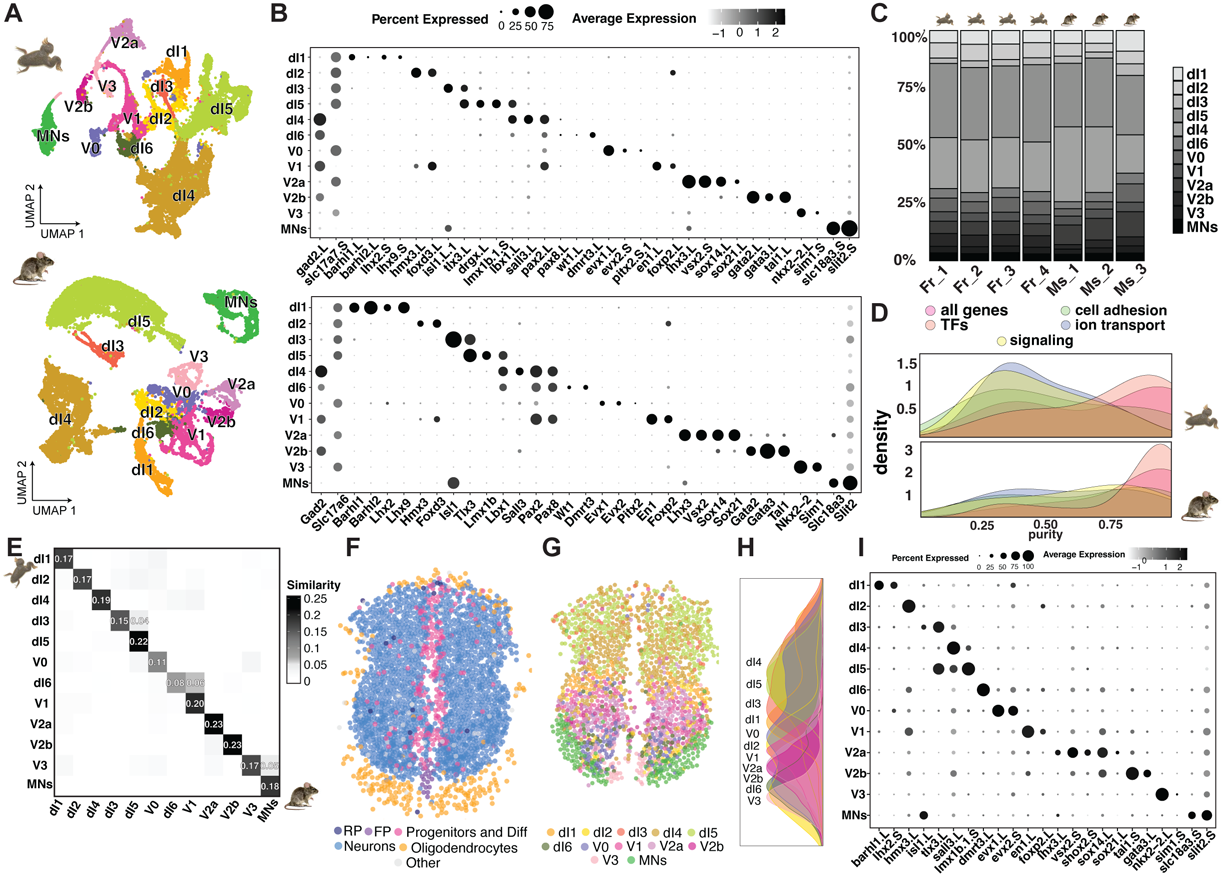
Conserved spinal neuronal architecture reveals shared developmental programs across evolutionarily distant tetrapods. **A.** UMAP representation of frog (top) and mouse (bottom) spinal neurons, color-coded according to their cardinal class identities. Each point corresponds to a single cell transcriptome, with cardinal class assignments determined by the expression of canonical markers. **B.** Dot plot showing conservation of cardinal class–defining gene programs across species. Orthologous cardinal class marker genes expressed in frog (top) and mouse (bottom). Dot size indicates the proportion of cells within each cluster expressing the marker, while color represents the average scaled expression level. **C.** Proportions of cardinal classes across frog (NF stage 54) and mouse (stage E12.5) replicates. F indicates frog and M indicates mouse, with numbers denoting biological replicates. **D.** Distributions of cardinal class purity, measured as the fraction of cells assigned to the dominant cardinal class, after re-clustering the data using different subsets of gene categories. For each subset, cardinal class identity preservation was assessed within the resulting clusters, enabling evaluation of how well specific gene categories capture major neuronal classes. Results are shown separately for frog (top) and mouse (bottom). **E.** Heatmap displaying pairwise similarity indices between frog and mouse cardinal classes, calculated from the integrated frog–mouse representation (Seurat; see **Methods** and **Fig. S3**). Heatmap represents the degree of transcriptional similarity between each frog and mouse cardinal class, providing a quantitative assessment of cross-species correspondence at the class level. **F.** Example of a NF54 frog spinal cord section with segmented cells labeled by coarse cell types, including roof plate, floor plate, progenitors, differentiating cells, neurons, oligodendrocytes, and other glial populations. **G.** Same section as in **F**, but displaying only neurons, which are color-coded according to their cardinal class identity. **H.** Density plot illustrating the scaled spatial distribution of cardinal classes along the dorsoventral axis, compiled across all frog spinal cord sections. **I.** Dot plot showing the expression of cardinal class markers in segmented and labeled cells across all frog spinal cord sections.

To investigate how gene categories contribute to cross-species segregation of cardinal classes, we performed clustering analysis using differential gene subsets, comparing transcription factors, cell adhesion molecules, and genes involved in ion transport and signaling. Following clustering, we quantified how well cardinal class assignments were preserved — calculating cluster purity by measuring the proportion of cells from the same cardinal class within each cluster. Cluster purity was highest when clustering with transcription factors or the full set of homologous genes, whereas other gene categories, such as those involved in adhesion or signaling, yielded lower or more variable purity scores (**Fig. 2D**). Transcription factors are thus the primary determinants of developing spinal architecture, not only within,^31^ but across evolutionarily distant species.

As an independent measure of the conservation of cardinal class identity across species, we integrated the mouse and frog datasets using multiple approaches (see **Methods**) and performed quantitative k-nearest neighbors (kNN) similarity analyses, quantifying the proportion of overlapping mutual nearest neighbors by species for each cardinal class pair. All methods revealed a high degree of one-to-one correspondence between cardinal classes in frog and mouse, validating the observed similarity in their overall expression profiles (k = 15, 20, 25; **Fig. 2E; Fig. S3**). Lastly, we examined transcriptional similarity between frog and mouse cardinal classes by calculating Pearson correlation coefficients for each cardinal class pair based on a subset of homologous gene expression profiles (see **Methods**; **Fig. S4**). Correlations were strongest within the same cardinal classes when using either all homologous genes or shared transcription factors (**Fig. S4A-B**). In contrast, gene sets related to other categories resulted in broader supergroups, with cardinal classes segregating by excitatory and inhibitory neuron identity (**Fig. S4C-E**). Nonetheless, some cardinal classes — including dI4, dI5, dI6 and motor neurons — retained their unique conserved code even for functional gene categories.

We also compared different stages of mouse development (E9.5 to E13.5) to our *Xenopus* NF54 single-cell dataset in terms of cardinal class composition (**Fig. S5**). Among these, the latest embryonic stage E12.5 in mice most closely resembled the overall structure of our developing frog neural dataset (**Fig. S5C**), consistent with our previous cross-species developmental alignment.^37^ We therefore focused on this stage and compared it to frog data to investigate whether even finer-scale neuronal subtypes — defined by previously characterized temporally regulated genes such as *nfia/nfib/neurod2/6* (N-division; late-born neurons) and *zfhx3/4* (Z-division, early-born neurons) — were also conserved within cardinal classes.^38^ By sub-clustering each cardinal class in both species, we found that this N/Z divisional structure was well conserved (**Fig. S6**). Thus, an ancestral, largely transcription factor–based architecture — spanning both cardinal class identity and temporal subdivisions — emerges as a conserved developmental feature of tetrapod motor circuits.

To validate and extend our single-cell RNA sequencing data to intact tissue, we conducted *in situ* spatial transcriptomic analysis of neuronal, cardinal and subcardinal molecular architecture using molecular cartography (Resolve).^39^ This method allowed us to probe the simultaneous expression of 100 targeted RNA sequences, chosen to span the defining molecular features of spinal neuron and cardinal class identity (**Table S1**), in intact tissue with nanometer resolution. Post-hoc cell segmentation of this data and further label transfer from our single-cell reference revealed the precise location of each cell and cardinal neuron type in the developing frog spinal cord (**Fig. 2F-H; Fig. S7-9**). Both coarse and finer scale distinctions between cell types were apparent: neurons, progenitors, oligodendrocytes, floor and roof plate cells, occupied distinct spatial domains (**Fig. 2F; Fig. S8**), and all cardinal classes were found *in situ* (**Fig. 2G; Fig. S9**), as in isolated single cells (**Fig. 2A-B**).

The mutually exclusive expression of core transcription factors again defined these classes, and as in mice^14^, each class occupied a differential dorsoventral position (**Fig. 2H-I; Fig. S9**). The dorsal spinal cord was predominantly composed of neurons from the dI4 and dI5 classes, the most prevalent among the dorsal neural populations. The ventral spinal cord displayed a characteristic dorsoventral layering of V0-V3, dI1-2, and dI6 interneurons, mirroring previous distributions in the mouse spinal cord^36^. Spatial transcriptomics data further revealed that the N- and Z-divisions segregate cardinal classes into lateral and medial populations as in mice (**Fig. S10**)^16^. Spatial mapping thus largely reflected the distributions observed in our single-cell data. These results together demonstrate high conservation of the cardinal class architecture at not only a molecular but also a spatial level between distant limbed vertebrates. This supports that a highly conserved developmental program specifies spinal neuron position, and by extension likely connectivity and function, across species.

### The spinal cord of adult frogs exhibits a diverse repertoire of cellular types

Given the striking similarity of developing spinal cord neural architecture between amphibians and mammals, yet distinct locomotor programs and sensory environments between species, we asked whether this developmental neural type similarity persists in adult stages. We hypothesized that adult neural types could emerge to support differences in motor and sensory repertoire between species that were absent or undetectable at earlier stages.

To test this hypothesis, we generated a complementary cell-type atlas of the adult amphibian spinal cord by isolating and sequencing single nuclei from sexually mature *Xenopus laevis* frogs (**Fig. 3A; Fig. S11**). Clustering of 28,264 single-nucleus transcriptomes from three biological replicates identified major cell types distinguished by established markers: *elavl4*, *nrxn3*, *slc17a7*, and *slit2* for neurons; *mbp* and *plp1* for oligodendrocytes; *mpz* for Schwann cells; and *slc4a4* for astrocytes (**Fig. 3B; Fig. S11D–G**). Notably, neurons were the most abundant population, comprising 11,831 nuclei (∼40%), and including spinal interneurons, motor neurons, and cerebrospinal fluid-contacting neurons (**Fig. 3C**). Initial low-resolution clustering of neurons revealed nine distinct neural groups (**Fig. S11H**): a *chAT*-expressing motor neuron cluster; three *sall3/pax2*-expressing inhibitory neuron clusters; four *lmx1b*-expressing excitatory clusters; and one large mixed cluster expressing *zfhx3/4* and markers of various cardinal classes (**Fig. S11I**). Drawing from published single-cell atlases in mouse^14,28^ and human^29,40^, we annotated *sall3/pax2*-positive clusters as dorsal inhibitory interneurons, *lmx1b*-positive clusters as dorsal excitatory interneurons, and the large *zfhx3/4*-positive mixed cluster as mid-ventral spinal interneuron populations (**Fig. 3D**).

**Fig. 3.**
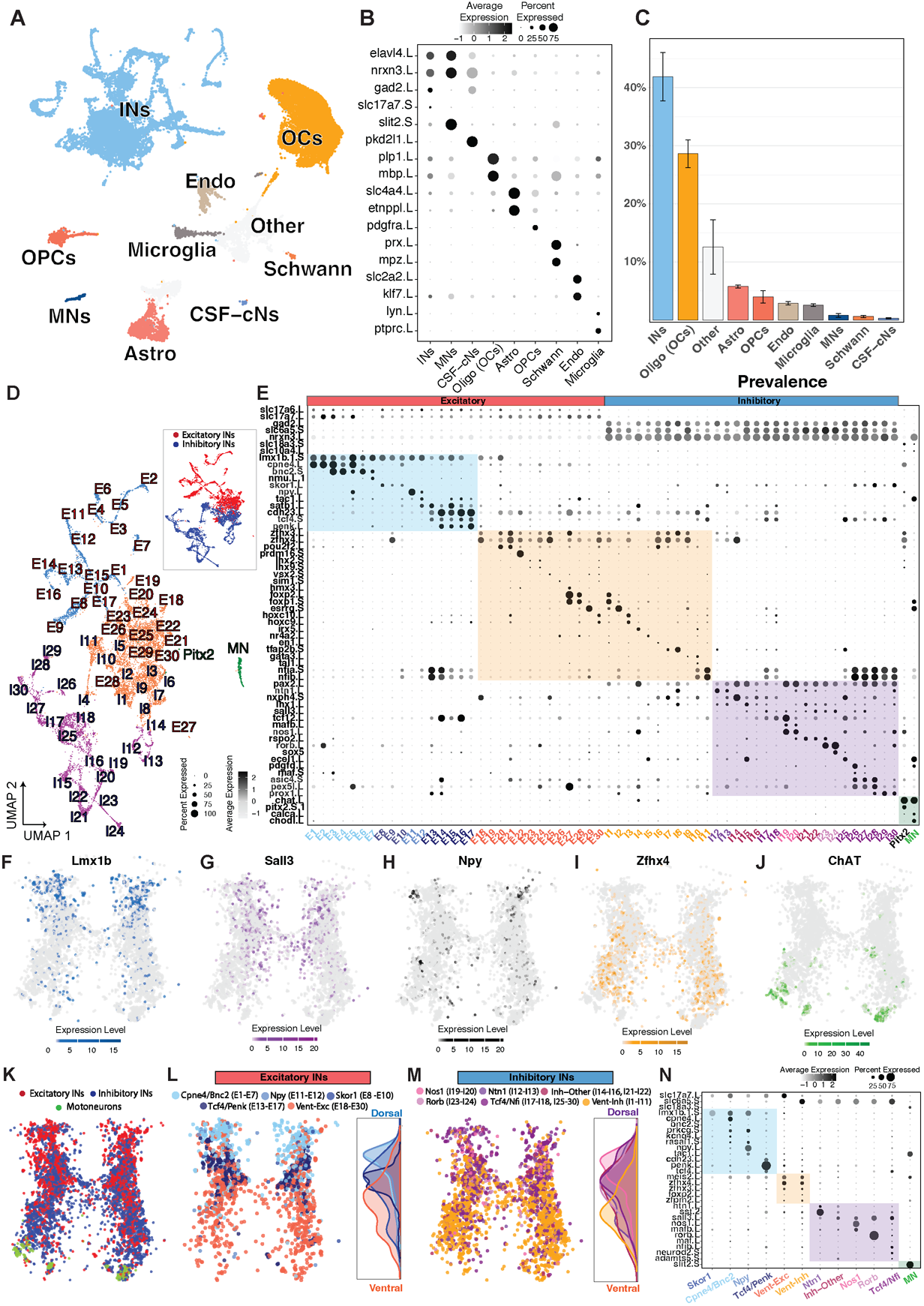
Single-cell and spatial atlas of the adult frog spinal cord. **A.** UMAP representation of color-coded by cell type: INs, interneurons; Oligo (OCs), oligodendrocytes; Endo, endothelial cells; Astro, astrocytes; MNs, motor neurons; OPCs, oligodendrocyte precursor cells; CSF-CN, cerebrospinal fluid–contacting neurons; microglia and Schwann cells; and “other,” indicating unclassified populations. **B.** Dot plot showing the expression of cell type–specific marker genes. Dot size reflects the proportion of cells within each population expressing the marker, while color intensity represents the average scaled expression level. **C.** Proportions of adult spinal cord cell types, with error bars representing the standard deviation across replicates. **D.** UMAP representation of frog neurons, color-coded by identity: light blue, dorsal excitatory; purple, dorsal inhibitory; green, motor neurons; and orange, ventral neurons. Inset, same neurons color-coded by excitatory (red) or inhibitory (blue) classification. **E.** Dot plot showing the expression of marker genes defining frog neural types. Dot size indicates the proportion of cells within each population expressing a given marker, while color intensity reflects the average scaled expression level. As in D, neurons are grouped by excitatory (red) and inhibitory (blue) identity, and further subdivided by dorsoventral location: light blue, dorsal excitatory; purple, dorsal inhibitory; orange, ventral neurons; green, motor neurons. **F-J.** Representative spinal cord section showing cell-level expression of marker genes: blue, *Lmx1b* (**F**); purple, *Sall3* (**G**); black, *Npy* (**H**); yellow, *Zfhx4* (**I**); and green, *ChAT* (**J**). **K.** Representative adult frog spinal cord section with segmented neurons labeled by neurotransmitter identity — excitatory (red) or inhibitory (blue) — or as motor neurons (green). **L-M.** Same spinal cord section as in **K** with either segmented excitatory neurons (**L**) or inhibitory neurons (**M**) labeled by neural type family. Inset on the right depicts the spatial distribution of each cell type aggregated across all sections.For **K-M**, labels were assigned by transfer from previously analyzed single-nucleus reference data. **N.** Dot plot showing the average expression of cell type marker genes in the spatial dataset, across segmented and labeled neurons from all analyzed sections. The plot includes cell types whose spatial distributions are shown in **L** and **M**, grouped by dorsal inhibitory, dorsal excitatory, ventral, or motor neuron identity.

Excitatory and inhibitory neurons of the dorsal and ventral spinal cord are classically thought to regulate sensory processing and motor output, respectively, and contain heterogeneous subpopulations in mammals.^30,41^ To understand whether these three coarse groups exhibited the same heterogeneity in frog, we further analyzed each group individually. Substantial intra-group diversity was found across all domains (**Fig. 3D-E**) with dorsal *lmx1b*+ excitatory and *sall3*+/*pax2+* inhibitory neurons the most heterogeneous, as in the mammalian adult spinal cord^14,29^. Within the *lmx1b*+ group, excitatory neuron families were marked by *cpne4*, *tac1*, and *tcf4*/*cdh23*/*penk*, aligning with Tac1^+^, Cpne4^+^, and Tcf4^+^ populations in mammals; similarly, *sall3/pax2*+ inhibitory neurons included subgroups expressing *nos1*, *rorb*, *maf*, and *ntn1*—with Nos1, Rorb, and Maf also observed in mammalian datasets.^14,15,42,43^ Presumed ventral neurons expressed *zfhx3/4*, *foxp2*, and a combination of cardinal markers including *lhx2/9*, *en1*, *gata3*, and *tal1*, consistent with ventral domain identities described in maturing mouse and human spinal cord.^10,13^

To evaluate whether the characteristic laminar distribution of adult neuronal populations in mammals is also found in amphibians, we performed high-resolution molecular cartography, paralleling our developmental analyses (**Fig. 2F-I, Table S2**). Following cell segmentation (see **Methods**), we transferred the labels from our adult *Xenopus* single-nucleus dataset to cell profiles on spinal cord sections. We confirmed that *lmx1b*-positive excitatory neurons (**Fig. 3F**) and *sall3*-positive inhibitory neurons (**Fig. 3G**), indeed localized to the dorsal spinal cord; *zfhx4*-positive neurons to the ventromedial (**Fig. 3I**) and *chAT*-positive motor neurons to the most ventral spinal cord (**Fig. 3J**). This agreed with the localization of similar, broad classes in mouse.^14^ We then extended this analysis to: (1) localize all inhibitory and excitatory interneurons, and motor neurons (**Fig. 3K; Fig. S12**) and (2) resolve finer-scale excitatory (**Fig. 3L; Fig. S13**) and inhibitory (**Fig. 3M; Fig. S14**) families as defined in the single-nucleus data (**Fig. 3D-E**). Among excitatory neurons, we identified classes with distinct laminar distributions: *npy*⁺ and *cpne4/bnc2*⁺ populations were localized to the superficial dorsal laminae; *skor1* and *tcf4/penk*⁺ populations were enriched in deeper dorsal layers; and the remaining excitatory populations to the ventral spinal cord (**Fig. 3L; Fig. S13)**. Within inhibitory neurons, we similarly delineated spatial distributions of defined classes: *nos1*⁺, *ntn1*⁺, *rorb*⁺, *tcf4/nfi*⁺ and other *sall3/pax2*⁺ inhibitory populations were predominantly localized to the dorsal region, with *nos1*⁺ cells most superficial; other inhibitory populations were concentrated within the ventromedial spinal cord (**Fig. 3M; Fig. S14)**. Lastly, we validated the accuracy of label transfer and the expression profiles for all spatially mapped populations, and found strong concordance of marker gene expression and cell type assignment between single-nucleus RNA-seq and spatial transcriptomic datasets (**Fig. 3N**).

Together, single-cell molecular and spatial profiling revealed a broad conservation of domain-level organization and selected interneuron families between amphibians and mammals. Despite this overall conservation however, the dorsal spinal cord exhibited notable species-specific differences: *npy* was expressed in dorsal excitatory neurons (**Fig. 3H; S13**), and *sox5* in dorsal inhibitory neurons, for example — whereas in mammals, these genes are restricted instead to inhibitory and excitatory neurons, respectively.^14^ To systematically resolve these evolutionary differences, we next undertook integrated analyses of adult spinal neuron datasets spanning a wider range of vertebrate species, including fish, frogs, mice, and humans.

### Dorsal excitatory neurons in mature spinal cord diverge in mammals

The dorsal spinal cord has long been posited to be the sensory processing center of mammals – integrating temperature, tactile and nociceptive signals to regulate pain, itch and touch motor reactions.^30^ Fish, frogs and other amphibians have a vastly different sensory environment than most terrestrial mammals: they are cold-blooded and often live at lower, more constant environmental temperatures; lack hairy skin; and have more limited tactile experiences.^44^ These ecological and physiological differences raise the possibility that dorsal spinal circuits — tasked with processing diverse sensory inputs — represent a focal point for evolutionary divergence across species. Our initial observations comparing adult frog and mouse spinal cord atlases support this hypothesis: while most adult frog neuron types retained coarse molecular features resembling those described in mammals, dorsal interneuron populations diverged in expression of several critical genes including Npy and Sox5 (**Fig. 3E**).

To resolve more comprehensively whether and how spinal neuron populations diverged across vertebrate evolution, we first quantitatively compared neuron types between the adult frog and published spinal cord datasets from both mouse^14,28^ and human^29,40^ **(Fig. 4A)**. Such a comparison between distantly related, but yet still four-limbed, vertebrates gave us the power to reveal species-specific features. As a first step in comparing spinal neuron composition across species, we utilized existing datasets from mouse and human (**Fig. S15**), and our frog data (**Fig. 3**), to evaluate spinal neuron distributions across species. Frogs showed a marked reduction in the proportion of dorsal excitatory neurons compared to mammals (**Fig. 4B**). Next, we used each species as the reference and transferred its neural-type annotations to the other two query species based on transcriptomic similarity (see **Methods**). This allowed us to measure how well neuron types defined in one species could be recovered in another. We found that label transfer accuracy was the highest between the two mammalian datasets, and consistently lower between either mammal and frogs (**Fig. 4C**), indicating substantial divergence of the amphibian spinal cord.

**Fig. 4.**
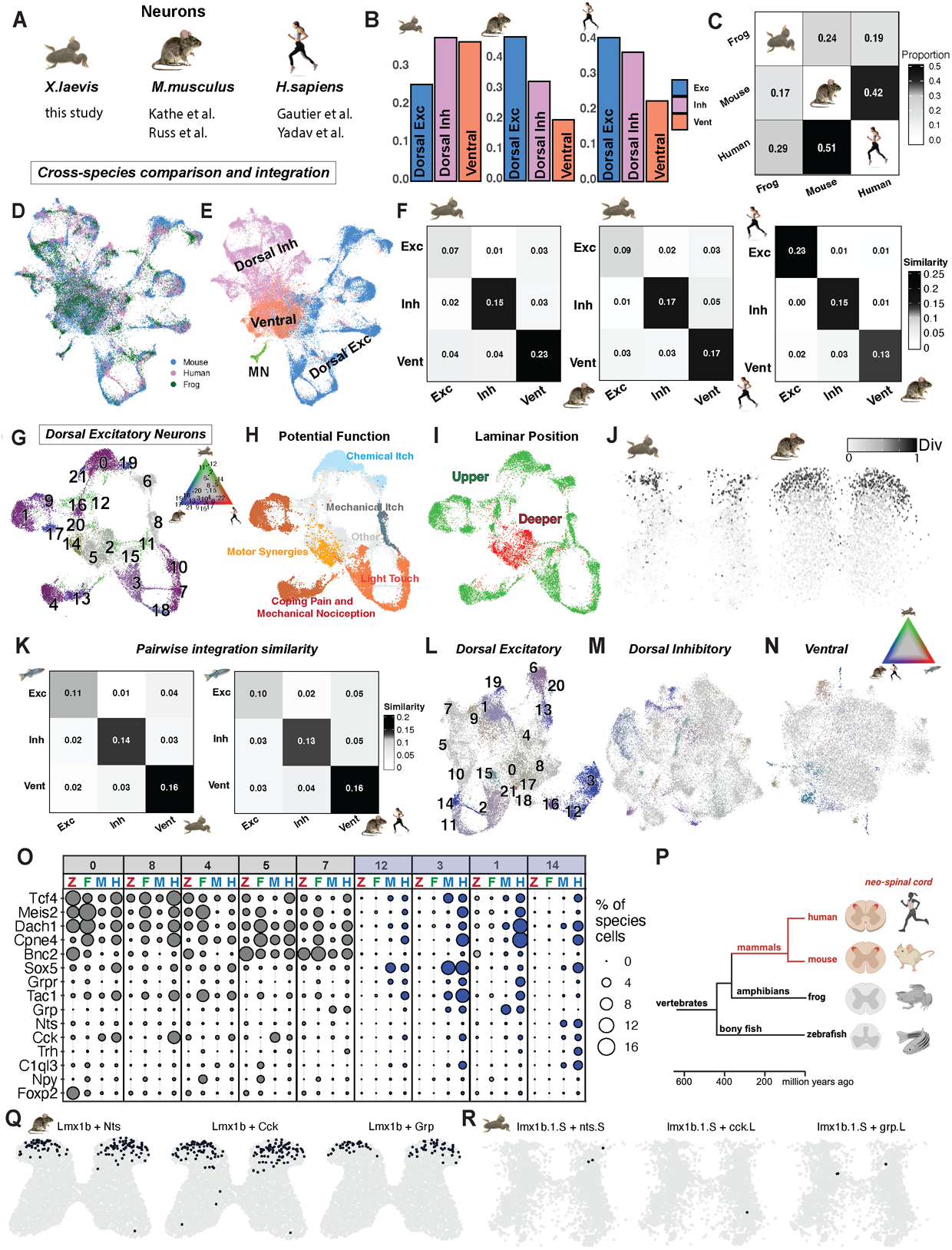
Selective divergence of dorsal excitatory neurons highlights evolutionary innovations in mammals. **A.** Metrics and references for the datasets used to compare spinal cord neural architecture across evolutionarily distant tetrapods (frog, mouse and human). **B.** Proportion of dorsal excitatory (Exc), inhibitory (Inh) and ventral (Vent) neurons in frogs, mice, and humans. **C.** Heatmap showing the proportion of cells in a query dataset that were successfully assigned labels from a reference dataset using label transfer (See **Methods**). **D-E.** UMAP-representation of frog, mouse and human CCA-integrated neural adult data labeled by species (**D**) and coarse neural type group (**E**). **F.** Heatmaps depicting pairwise similarity indices among coarse neural type groups within the integrated frog–mouse–human dataset, with each panel representing a comparison between two species. **G–I.** Integrated UMAP representation of frog, mouse, and human dorsal excitatory neurons, color-coded by species mixing (**G**), candidate literature-derived functions (**H**; only mouse cells labeled), and laminar position (**I**). **J.** Representative spinal cord sections from frog (left) and mouse Visium (right) spatial datasets, color-coded by conservation or divergence score. Spatial cells were annotated with cell type identity through label transfer from reference cells classified as conserved or divergent, respectively. **K.** Heatmap depicts the pairwise similarity scores of dorsal excitatory, dorsal inhibitory and ventral populations from zebrafish versus frogs (left) and zebrafish versus mammals (right). **L-N.** Integrated UMAP representations of zebrafish, frog, mouse, and human neurons for dorsal excitatory (**L**), dorsal inhibitory (**M**), and ventral (**N**) neurons, color-coded by species-mixing values. Inset depicts mixing values across populations with red color representing zebrafish, blue mammals (mouse and human), and green - frog. **O.** Dotplot showing the expression of dorsal excitatory neural type markers within integrated clusters of zebrafish, frog, mouse and human data. The color of the dot represents the value of species mixing for that specific cluster. Dot diameter shows the percentage of cells expressing this marker within all neurons from that species. **P.** Schematic depicting evolutionary innovation of dorsal excitatory neurons, the neo spinal cord (red), in mammals. **Q-R.** Validation of mammalian-specific dorsal excitatory subtypes via spatial transcriptomics. Number and position of Lmx1b-positive dorsal excitatory neurons expressing Nts, Cck and Grp in adult mouse (**Q**) and frog (**R**) spinal cord.

To further investigate this species divergence in a shared molecular framework, we integrated transcriptomic data from frogs, mice, and humans using multiple methods (see **Methods**). In this integrated space, dorsal excitatory neurons from frogs aligned poorly with their mammalian counterparts, whereas dorsal inhibitory and ventral neurons showed more consistent overlap (**Fig. 4D–E; Fig. S16**). To capture this difference quantitatively, we applied a kNN-based similarity metric to all pairwise species comparisons (see **Methods,** as in **Fig. S3**). For the dorsal excitatory population, the similarity score was consistently lower between frog-to-mammal than human-to-mouse comparisons (**Fig. 4F**). This dorsal divergence was not seen at earlier developmental timepoints (**Fig. S17**), supporting that these mammalian-specific evolutionary divergent populations emerge later in development.

As an independent method to validate dorsal divergence between species, we re-clustered dorsal excitatory, dorsal inhibitory, and ventral neurons and calculated species mixing scores for each resulting cluster (**Fig. 4G; Fig. S18**; see **Methods**). Based on these scores, subpopulations were defined as either conserved or species-specific. This analysis revealed numerous dorsal excitatory clusters composed predominantly of either mammalian-or frog-specific neurons (**Fig. 4G**). Among the mammalian-specific clusters, most contained both mouse and human cells, indicating a shared mammalian rather than a species-restricted subtype framework. In contrast, for dorsal inhibitory and ventral neurons, clusters showed substantial species mixing, and stronger conservation of subtype identity between amphibians and mammals (**Fig. S18**). Our results thus suggest that dorsal excitatory neurons have undergone greater subtype level divergence than inhibitory or ventral populations.

In mice, excitatory neurons exhibit distinct laminar distributions. Deeper neurons are associated with motor coordination and synergies, whereas superficial neurons connect with sensory afferents encoding pain, itch, temperature, and pleasant touch.^45–48^ To explore functional divergence across species, we mapped these previously annotated categories of mouse excitatory neurons onto our integrated multi-species dataset. We found that neurons implicated in pain modulation and light touch predominantly fell into mammalian-specific clusters, along with a subset of neurons linked to chemical itch **(Fig. 4H**). In contrast, neuronal populations associated with mechanical itch, the motor-related deep dorsal horn, and chemical itch were part of conserved clusters. Gene expression analysis revealed key differences in molecular signatures. For example, *Nts*-positive dorsal excitatory neurons were mammalian-specific (**Fig. S18D, F**), a population previously linked to pain sensitization and corrective tactile reflexes,^49,50^ and in frog but not mouse or human, *Npy* was expressed in dorsal excitatory neurons, suggesting divergence in nociceptive circuits **(Fig. S18E, F**). Unlike their dorsal excitatory counterparts, dorsal inhibitory and ventral interneurons were less divergent, with most populations conserved but the Rorb-/Pdyn-positive dorsal inhibitory population, also involved in mechanical pain and itch,^51^ being mammalian-enriched (**Fig. S18G-I**) and all major ventral classes well represented in all species (**Fig. S18J**). As expected, these functional distinctions also correlated with spatial organization.^50^ Transcriptomically-defined, species-specific populations mapped to known upper lamina in mice (**Fig. 4I**), suggesting evolutionary specialization of the superficial spinal cord. Further supporting this, we projected conserved and divergent populations from each species onto existing spatial data (see **Methods**). Populations identified as divergent based on integrated clustering consistently mapped to the most superficial lamina of the dorsal spinal cord (**Fig. 4J**). In mice, this region is tightly associated with sensory signal processing and integration.^30^

### Dorsal divergence of spinal cord cell types persists from fish to humans

To determine whether dorsal spinal cord divergence reflected a broader evolutionary pattern across vertebrates, we re-analyzed publicly available single-cell data^52^ to characterize the neural architecture of the adult zebrafish spinal cord. We clustered zebrafish neurons and performed label transfer from mouse and frog to identify major neural domains — dorsal excitatory, dorsal inhibitory, and ventral neurons (**Fig. S19E,F).** As in tetrapods, zebrafish neurons were organized into groups, expressing conserved markers such as Lmx1b for dorsal excitatory neurons, Pax2 for dorsal inhibitory neurons, and canonical ventral markers (**Fig. S19G,H,M,N**). Within each group, we identified finer-grained subpopulations, including those with species-conserved expression profiles — such as Bnc2, Maf, and Nmu within the dorsal excitatory domain, or Lhx1 and Nos1 within the dorsal inhibitory domain^14,15^ — as well as zebrafish-specific transcriptional signatures not present in tetrapods, including a population of Foxp2/Lmx1b-positive neurons (**Fig. S19N**). These findings demonstrate a largely conserved vertebrate organization of the adult spinal cord extending to teleosts.

Given this conserved coarse organization of zebrafish spinal cord cell types, we then asked whether the pronounced divergence of dorsal excitatory types was a lineage-specific feature or part of a broader vertebrate trend. To address this, we performed pairwise integration of zebrafish neurons with neuronal data from either frogs or mammals. In this and further analyses, mouse and human were grouped as a single ‘mammalian’ species due to their high degree of molecular similarity observed previously. Between these evolutionarily distant clades, again the most prominent decrease in molecular similarity was observed for dorsal excitatory neurons in both pairs (**Fig. 4K**). Cross-species integration of fish, frogs and mammals provided further support for selective divergence of the dorsal excitatory domain compared to other groups of neurons (**Fig. 4L-N**).

With four species in our analysis, we sought to define the molecular nature of dorsal neuron divergence. We evaluated gene expression profiles across integrated clusters for dorsal excitatory neurons (see **Methods**). Conserved excitatory populations included neurons expressing Tcf4/Meis/Dach1, markers of the deep dorsal horn in tetrapods (**Fig. 3**), and Bnc2/Cpne4, a well-established dorsal population associated with chemical and mechanical itch in mice (**Fig. 4O**). Divergent populations were defined by species-specific gene expression patterns. The most prominent of these included Sox5-positive mammalian-specific neuronal types, such as Grpr-, Tac1-, Grp- and Nts-expressing populations, which are implicated in pain modulation and localize to superficial spinal laminae. Independent spatial transcriptomic analysis validated the mammalian-specificity of these expression patterns (**Fig. 4Q; Fig. S20**). Together, these results demonstrate a key principle of spinal cord evolution: dorsal excitatory neurons, key for sensory integration and modulation, show the greatest molecular divergence across vertebrates and contain mammalian-specific subpopulations that comprise a putative, recently evolved neo-spinal cord.

## Discussion

In this study, we generated spatially resolved cell type atlases of the amphibian spinal cord at both developing and adult stages. Comparing these atlases to data from zebrafish, mice and humans, we trace the origins and evolution of the vertebrate spinal cord from teleosts to mammals. During development, we discover near-perfect conservation of the cardinal cell type architecture of the developing spinal cord. By contrast, at adult stages, excitatory dorsal neurons have selectively diverged in mammals.

Conservation of the hallmark cardinal class architecture of the developing spinal cord between amphibians and mammals, which last shared a common ancestor nearly 360 million years ago, argues for a deeply conserved, developmental blueprint of cell-type specification programs. As cardinal identity in mammals is critical to define a neuron’s functional properties, by extension, this conservation also argues for a similar functional organization of motor circuits across species. As in mice, each cardinal class in other vertebrates would thereby modulate a specific motor feature: V2a interneurons driving motor output by rhythmically exciting motor neurons; V0 inhibitory neurons coordinating left-right movement; inhibitory dI6 neurons regulating speed transitions and gaits; and V1 neurons the alternation of flexor-extensor muscles and the in-phase inhibition of motor neurons.^1^ This hypothesis that the same core cardinal building blocks are used across species, despite the wide range of vertebrate movement, is supported by functional perturbations in zebrafish^26,53–56^ and more recently, initial studies of interneuron function in frogs.^37^ Our study supports that such functional blocks originate from deeply conserved, developmentally specified molecular units. Such conservation points to other potential sources of species-divergent motor patterns, including variant molecular, connectivity, or physiological profiles of cardinal subtypes or supraspinal modulation of these conserved units.

This conservation also raises the possibility that such spinal blueprints may span even larger evolutionary distances, perhaps even beyond vertebrates. With the highly conserved, molecular signatures we identify, it is now possible to probe their evolutionary depth. In the nerve cord of the ascidian *Ciona intestinalis*, considered an evolutionary precursor to the vertebrate spinal cord, evidence of homologous transcription factors support the early emergence of cardinal classes.^57^ Even in annelids, there is initial evidence of nerve cord progenitor domains similar to those of vertebrates.^58^ Further research is needed to assess the ubiquity of neural type homology across bilaterians, including annelids, amphioxus, ascidians and beyond. Such analysis will reveal whether a common blueprint is necessary for movement in both non-vertebrate and vertebrate species.

Our cross-species analysis also demonstrates that the molecular nature of this conserved developmental organization largely hinges on transcription factors, raising the importance of investigating their downstream functional gene targets. Moreover, we observe that the temporal patterning factors of mammalian spinal cord,^38^ which play a critical role in determining neuronal position and projection range^16^, also are shared between amphibians and mammals, further dividing each cardinal class into non-overlapping, molecularly-distinct subpopulations. This demonstrates that the basic mechanisms specifying other non-cardinal aspects of neural diversity in the spinal cord are also highly conserved across vertebrates, and supports that extensive spinal neuron subtype heterogeneity is not just a mammalian-specific feature but rather a common characteristic of the vertebrate spinal cord.

The remarkable similarities in neural architecture during development however sharply contrast with the divergence between species at adult stages. This immediately begs the question of how a conserved blueprint develops into a divergent endpoint. This question of how cell types change within and across species from the developing to mature stage has recently been an active area of study. Developmental conservation and adult divergence is also observed in the brain. In the pallium, despite high conservation during embryonic development, neural types diverge at adult stages.^59^ Mechanistically, this adult onset divergence could be the result of sequential birth timing, with the most divergent populations, the last born. Recent studies indicate that the most superficial dorsal horn types, the most divergent at adult stages, are indeed later born.^15^ This raises the attractive mechanistic possibility that evolutionary innovation may have selectively acted on the last cell divisions to drive species-specific cellular types, a hypothesis that too is gaining support in the brain.^60^ Alternatively, the initial formation of neural circuits via highly conserved transcriptional programs could be followed by a second period of species-specific cell-type differentiation, in which functional gene categories that are unique to each species are now upregulated.

Notably, our comparisons at adult stages localize this divergence in excitatory neural types to the most superficial regions of the dorsal spinal cord. This selective divergence of excitatory, as compared to inhibitory, neuron types echoes recent findings in the brain across species,^61^ implying evolutionary flexibility of excitatory circuits is a common feature of nervous systems. In mice, such dorsal interneurons are largely associated with sensory integration and processing.^30^ This contrasts with the ventral, mid and deep dorsal horn regions of the spinal cord, which are primarily associated with motor execution.^45^ Such selective divergence of the most superficial dorsal spinal cord was most likely thus an innovation of sensory processing, either directly via peripheral sensory neurons or indirectly via spinal cord-to-brain interconnectivity. The mammalian-specific cell types we identify, including those positive for Grpr, Trh, and Nts have a well-established role in nociception in mice,^62^ potentially highlighting evolutionary changes in pain sensation and processing. Interestingly, a significant portion of mammalian-specific clusters could also be linked to populations activated in mice during sensory stimulation but not during walking.^63^ Recent analysis of sensory neurons across species, including amphibians, argues for a coarse conservation of sensory types but notable differences in their receptor expression.^64^

An additional evolutionary observation emerging from our study is that the diversification of dorsal neural types across evolution correlated with the reappropriation of neuropeptides. Exemplifying this, neuropeptides such as *Npy,* predominantly expressed in inhibitory neurons in mice, are expressed in excitatory neurons in frogs. Other neuropeptides like Cck and Nts are species-specific in their expression: in the mouse and human but largely absent from the frog spinal cord. Recent comparisons between mice, macaque, and humans also reveal slight differences in neuropeptide gene and receptor expression in the spinal cord,^65^ albeit less numerous than those between amphibians and mammals. These results align with growing evidence that evolution of neuropeptides and their cell-type-specific expression is a crucial mechanism for endowing a neural type with a unique expression and functional landscape.^66–68^

In light of recent comparisons across vertebrates that highlight extensive neural type diversification in the brain during evolution,^69,70^ and previous evidence of significant differences in brain-to-spinal cord projection across vertebrates,^5^ including the absence of direct dorsal telencephalon-to-spinal projections in amphibians, the differences we observe could also reflect more recently evolved descending and ascending pathways. The superficial laminae of the dorsal cord, which exhibit the most mammalian-specific subtypes, are known to project to the brain.^71^ An increase in brain-to-spinal cord connectivity may thus have accompanied the evolutionary expansion of the forebrain, exemplified by its increased size, cell number and subtype diversity in mammals.^72^ Further studies visualizing brain-spinal cord circuits in lesser studied, more basal vertebrates using recently developed tools^73^ will reveal how mammalian-specific pathways evolved and relate to the species-specific subtypes identified here.

Our work examining spinal cord cell types across nearly 500 million years of vertebrate evolution thus provides some of the first insights into the evolutionary origins of spinal cell types, complementing recent cross-species analyses of the vertebrate brain. A common theme of nervous system evolution emerges from fish to mammals: a shared developmental blueprint of transcriptionally defined cell types exists but this blueprint diverges at adult stages, when selective radiation of excitatory neuron types becomes evident in mammals. This divergence maps onto the superficial dorsal layers of the adult spinal cord, making the prediction that higher-level sensory processing is a recently evolved mammalian trait of the “neo-spinal cord.”

## Methods

### Animals

To generate *Xenopus laevis* spinal neuron single-cell data at post-metamorphic NF stage 54 froglets, we used transgenic *Xla.Tg(tubb2b:maptGFP)Amaya* (EXRC), expressing GFP under pan-neuronal *tubb2b* transcriptional control, enabling us to enrich for neuronal cells. To generate adult Xenopus laevis spinal neuron nuclei data, pigmented wildtype animals were used. All animals were bred and raised at the Institute of Science and Technology Austria (ISTA) following protocols approved by the local authorities (permit number: 2020-0.550.806). Tadpoles of both sexes were used indiscriminately, as sex cannot be determined at this early stage. Both sexually mature males and females were used for adult experiments.

### Spinal Cord Dissociation and Cells/Nuclei Isolation

For NF stage 54 developing froglets, spinal cords were isolated from ten transgenic animals and pooled into 4 technical replicates. Acquired tissue was dissociated with liberase^74^ to obtain single-cell suspension according to previously published methods, and FACS-sorted for the presence of GFP using a BD FACS Aria III Cell Sorter.

For adult *Xenopus laevis*, we prepared single-nuclei instead of single cells, as adult neurons can be sensitive to single-cell dissociation procedures. Two male and one female adult spinal cords were isolated, flash frozen, and nuclei were extracted separately for each animal, totalling three biological replicates, according to established detergent protocols for adult mouse spinal cord nuclei isolation.^75^

### 10X Genomics Sample Processing - Library Preparation and Sequencing

Cells/nuclei were then processed following the Chromium Single Cell 3’ v3.1 protocol and Chromium Reagent Kits were used to prepare single-cell libraries (10X Genomics). Sequencing was performed on a NovaSeq (Illumina).

### scRNA- and snRNA-seq Data Processing and QC

10X CellRanger v5.0.0 was used for gene alignment to obtain gene-barcode matrices. The genome used for the reference was Xenla 9.2 (X. laevis v9). Then, individual count matrices were imported into R (v4.4.2), converted into Seurat data object, and further analyzed with Seurat v5.

For developmental *Xenopus* data, raw count matrices were processed to remove low-quality cells and doublets. To exclude low-quality or dying cells, we assessed multiple quality features, including the number of detected genes, total read counts per cell, and the proportion of mitochondrial transcripts. Rather than applying manually defined thresholds, we modeled the distribution of these features using a binomial mixture framework implemented in the *mixtools* package in R.^76^ Thresholds were then selected based on the inferred mixture components, enabling automated separation of low-quality and viable cells. Doublets were identified and removed using the *DoubletFinder* package in R.^77^

### Clustering and Cell Type Assignment – General Overview

To cluster cells and define cell populations in the data we use the Seurat package in R.^78^ To perform principal component analysis we use the RunPCA function on the basis of the most variable features in the data that are defined with the function *FindVariableFeatures*. Then, we construct k-nearest neighbor (KNN) graphs with function *FindNeighbors* on the basis of the defined principle components and perform hierarchical clustering to group cells by their transcriptional proximity with *FindClusters* function based on the KNN graph. We assign cell-type labels and identify populations on the basis of already known spinal cord cell types markers and/or differentially expressed genes for individual clusters that are calculated with *FindMarkers* function in *Seurat*. We visualize cell populations and gene expression data with non-linear dimensional reduction method UMAP plots that use principal components as an input (*Seurat*).

### Clustering and Neural Type Assignment of Xenopus laevis data

For frog developmental data, raw counts were processed in R (v4.x) with Seurat (v5). After QC (see “scRNA- and snRNA-seq Data Processing and QC”), neuronal cells were isolated based on canonical markers (e.g., elavl4.L, snap25.L) and used for downstream analysis. Data were log-normalized, highly variable features were selected, and PCA was computed (50 PCs). We built a k-nearest-neighbor (KNN) graph on PCA space (dims used for neighbors/UMAP stated below) and performed Louvain clustering at high resolution to separate partially overlapping cardinal classes. Neuronal identities were assigned by combined marker patterns (examples: dI1—*lhx2/9.S*, *barhl1/2.L*; dI2 - hmx3.L, foxd3.L; dI3—*tlx3/drgx.L*; dI5— lmx1b1.S; dI6—wt1/dmrt3.L; V0—evx1.L, pitx2.s; V1—en1/foxp2; etc). Ambiguous or mixed-identity clusters were iteratively subclustered at higher resolution and relabeled using differential expression (Seurat FindMarkers, min.pct = 0.25, logfc.threshold = 0.25) and diagnostic dot/feature plots. Low-quality clusters detected post hoc (e.g., very low gene counts, nonspecific signal) were removed and the graph was rebuilt before a final clustering pass.

For frog adult data, data from three spinal cords / replicates were processed separately with SCTransform (variable.features.n = 7000) and integrated using Seurat’s SCT workflow (SelectIntegrationFeatures, PrepSCTIntegration, FindIntegrationAnchors, IntegrateData). Dimensionality reduction was performed with PCA (50 components) and UMAP (dims 1–50). A k-nearest neighbor (KNN) graph was built on these dimensions, and cells were clustered with FindClusters at resolutions ∼0.5–1.5 across iterative steps. Major cell classes were assigned by canonical markers — excitatory neurons (Slc17a7), inhibitory neurons (Gad2), motor/cholinergic neurons (Slc18a3, Pitx2), oligodendrocytes (Plp1, Mbp), astrocytes (Slc4a4), OPCs (Pdgfra), microglia (Ptprc), endothelial cells, and CSF-contacting neurons (Pkd2l1). Non-neuronal populations were excluded from subsequent neuronal refinement. For neuronal subtyping, we subset neurons and repeated the SCT integration across replicates, followed by PCA (50 PCs), KNN construction, UMAP, and re-clustering at higher granularity. A small low-quality subset within one cluster was identified by re-clustering and marker inspection and removed, after which we re-ran PCA/UMAP/FindNeighbors/FindClusters on the cleaned neuron set. Mixed-identity clusters that combined excitatory and inhibitory signatures were explicitly subclustered at higher resolution and split into excitatory and inhibitory neurons. We then organized neurons into three groups — dorsal excitatory (lmx1b.L), dorsal inhibitory (pax2.L, sall3.L), and ventral/mid (e.g., zfhx3/4.L) based on low resolution clustering (resolution = 0.07) — and carried out targeted reclustering within each group to obtain fine-grained neural identities. Neural group identity was further confirmed with spatial transcriptomics (See ***Spatial Transcriptomics - Label Transfer and Visualization***)

### Re-analysis of Public Data Sets from Mouse, Human and Zebrafish and Coarse Neural Type Assignments

We re-analyzed the following datasets: mouse developmental (e9.5-e13.5) scRNA-seq data from Delile et al.; mouse post-natal scRNA-seq and spatial (Visium) data from Russ et al. and Kathe et al.; human adult scRNA-seq data from Yadav et al. and Gautier et al.; and zebrafish adult scRNA-seq data from Saraswathy et al. For mouse developmental data, we re-analyzed embryonic mouse spinal cord scRNA-seq data (E9.5–E13.5) from Delile et al. (2019) [https://github.com/juliendelile/MouseSpinalCordAtlas], which provides raw UMI count matrices and accompanying annotated metadata (stages and replicates). Raw counts were processed in R (v4.x) using Seurat (v5). Ensembl gene IDs were converted to gene symbols via BioMart; ambiguous mappings (multiple Ensembl IDs mapping to the same gene) were retained as Ensembl IDs, while the remainder were converted to corresponding gene names. Cells were filtered to remove low-quality barcodes (nFeature_RNA ≤ 1000, **S2A**).

Data were log-normalized, highly variable features identified, and dimensionality reduction performed with PCA (50 PCs computed). To reduce technical signal associated with cell-cycle and ribosomal programs, downstream analyses used PCs 5–30. UMAP embeddings were computed on these dimensions, followed by graph-based clustering. Neuronal populations were identified based on canonical marker expression (e.g., Elavl4, Snap25, Slc18a3, Isl1/2, Lmx1a, Pax6) and retained for downstream analysis (**S2BC**). Cells were split by replicate and integrated with Seurat’s workflow with prior SCTransform normalization per replicate to mitigate batch effects (FindIntegrationAnchors with normalization.method = “SCT”; IntegrateData with normalization.method = “SCT”, **S2DE**). After integration, clustering was repeated (resolution = 2.0), yielding discrete neuronal clusters (**S2FG**). Neuronal progenitor contamination was removed (**S2F**) and neuronal clusters were annotated into cardinal classes (dI1–dI6, V0–V3, motor neurons) using established markers (Lhx2/9, Barhl1/2, Isl1, Lhx3, Vsx2, En1, Evx1/2, Pitx2, Nkx2-2, Sim1, Slc17a6, Gad2, Pax2, etc., **S2G**). In one mixed cluster, subclustering revealed distinct V0 (Pitx2, Evx1/2) and V1 (En1, Foxp2) populations, which were subsequently relabeled. The final processed object contained only neuronal cells assigned to canonical cardinal classes and was used for cross-species comparisons.

For Russ et al. mouse post-natal snRNA-seq data, raw count matrices were downloaded from GEO [GSE158380]. In addition, we obtained the authors’ integrated reference Seurat object (assembled from previously published postnatal datasets) and their accompanying neural-type annotations, which were used for label transfer (see below). Counts were processed: log-normalization, variable feature selection, PCA (50 PCs), UMAP, and graph-based clustering. Neuronal populations in the postnatal dataset were annotated by label transfer from the integrated reference; predictions were filtered at a score threshold of 0.5 to retain high-confidence assignments, and cells annotated as neurons or motor neurons were retained and reprocessed. Clusters corresponding to CSF-contacting neurons (marked by Pkd2l1) were excluded to focus on dorsal/ventral interneurons and motor neurons. The resulting object comprised high-quality neuronal populations annotated according to reference-derived neuronal subtypes and was used in downstream analyses (**S15E**).

For Kathe et al. adult mouse snRNA-seq data, raw count matrices, feature annotations, and accompanying metadata were obtained from GEO [GSE184370]. Cells annotated as neurons in the provided metadata were subset and reprocessed in R (v4.x) using Seurat (v5): log-normalization, variable feature selection, PCA (50 PCs), UMAP, and graph-based clustering. Clusters corresponding to CSF-contacting neurons were identified and removed. The final processed object consisted of neuronal populations (interneurons and motor neurons).

For Gautier et al. adult human spinal cord snRNA-seq data, the authors provided a Seurat object containing neuronal populations, which we downloaded directly from GEO [GSE228778]. To ensure consistency with other datasets, neuronal cells were reprocessed in R (v4.x) using Seurat (v5): raw counts were extracted, normalized, highly variable features identified, and dimensionality reduction performed with PCA (50 PCs) followed by UMAP embeddings.

For Yadav et al. adult human spinal cord spinal cord snRNA-seq data, aggregated post-QC count matrix and accompanying metadata were downloaded from GEO [GSE190442] and imported into Seurat (v5). Gene symbols were standardized, and cells were processed with log-normalization, variable-feature selection, PCA (50 PCs), UMAP, and graph-based clustering. Neurons were subset using the authors’ provided annotations and further reclustered. Low-quality cells were removed.

Coarse neural group identities — dorsal excitatory (Exc), dorsal inhibitory (Inh), ventral/midline (Vent) (and MN where applicable) — were assigned to the four mammalian datasets (mouse: Russ, Kathe; human: Gautier, Yadav) by Seurat label transfer from the integrated Russ et al. mouse reference. We first collapsed the Russ fine labels (“Excit-xx”, “Inhib-xx”, “MN-α/γ”, “PGC”) into these coarse domains according to their laminar/topographic placement as summarized in the study, then transferred labels with FindTransferAnchors (normalization.method = “LogNormalize”, reduction = “pcaproject”, features = intersect(genes)) followed by TransferData. Coarse calls were taken with GetTransferPredictions(score.filter = 0.2) to maximize recall, while fine Russ subtypes were optionally transferred with score.filter = 0.5 for higher precision. For all queries we set DefaultAssay = “RNA”. In Gautier, a small low-resolution cluster (resolution = 0.05) with inconsistent scores was reassigned to Exc before finalizing coarse labels. Cells below the score thresholds remained unassigned and were excluded from summaries. In Yadav these assignments were used to guide renaming of their provided neural type labels into these coarse groups (**S15A-D**). We also applied label transfer from the integrated Russ et al. reference to our adult frog dataset, which resulted in the reassignment of populations E18– E20 from mid/ventral to dorsal excitatory, and E23–E24 as dorsal excitatory and mid/ventral, respectively.

To then compare the proportions of coarse neuronal classes (dorsal excitatory, dorsal inhibitory, and mid/ventral) across adult datasets, we used our frog data, the dataset of Kathe et al. for mouse, and the dataset of Gautier et al. for human, respectively.

For Saraswathy et al. adult zebrafish spinal cord snRNA-seq data, raw count matrices, barcodes, and feature files were obtained from GEO [GSE235395]. Ambient RNA was corrected using DecontX, and data from multiple samples were subsequently merged into a single Seurat object and clustered. Neuronal populations were identified using canonical markers (*snap25a, elavl3*), subset, and transformed into mouse gene space (see **Cross-species ortholog mapping**) to enable identification of coarse neural domains via reference-based transfer from frog and mouse datasets. The neuronal subset was then re-clustered (**Fig. S19A–C**). Residual non-neuronal contamination was identified and removed (**Fig. S19D**). Initial clustering yielded 25 clusters, which were classified into dorsal excitatory, dorsal inhibitory, and ventral populations based on label transfer results and marker analysis (e.g., *Lmx1b* for dorsal excitatory, *Pax2* for dorsal inhibitory, *Meis2* for ventral), as well as motor neurons. Contaminant clusters were identified such as sensory neurons (*Prph*), injury-responsive neurons (*Atf3*) (**Fig. S19I–J**), and an oligodendrocyte-enriched cluster identified during ventral re-clustering and all were further excluded from the analysis (*Plp1, Mpz* expression, absence of neuronal markers; **Fig. S19K–L**). Final neuronal annotations were derived from the remaining high-confidence neuronal clusters, and coarse identities (dorsal excitatory, dorsal inhibitory, mid/ventral) were assigned according to marker gene expression and label transfer results (**Fig. S19M-N**).

All the resulting objects were used for downstream cross-species comparisons.

### Cross-species Ortholog Mapping

To enable cross-species integration, frog and zebrafish datasets were transformed into a mouse gene space using gene correspondence tables. For *Xenopus laevis*, isoforms from both long (.L) and short (.S) genomes were aligned to mouse genes using a HMMER3-based protein-homology pipeline (**Table S3**). Counts from isoforms mapping to the same *Mus musculus* gene were summed, yielding a frog gene expression matrix in mouse gene format. For zebrafish, orthology relationships were obtained using OrthoFinder (**Table S4**). As with frog, counts from multiple zebrafish genes mapping to the same mouse gene were collapsed by summation, and genes lacking an ortholog were discarded. In cases of one-to-many mappings, the first listed mouse ortholog in the correspondence table was retained. Human data were used without modification due to the consistent gene nomenclature with mouse data.

### Cross-species Neural Type Mapping and Label Transfer

We quantified the *proportion of successfully mapped cells* as the proportion of cells in a query dataset that could be successfully assigned labels from a reference, calculated as the number of labeled cells divided by the total number of query cells. To achieve this, we performed pairwise label transfer between adult datasets (frog, mouse - Kathe et al., human - Gautier et al.) as follows:

1. Each dataset was processed with SCTransform (Seurat v5), regressing technical covariates as appropriate (e.g., library size/feature counts and dataset-specific batch/sample terms), and variable features were selected (10,000 per dataset). To ensure a shared feature space, we intersected the variable features across datasets and computed PCs for each dataset on this intersected shared set which were further used for clustering.
2. To select reference cell identities to be mapped, we use our curated neuronal type labels for the frog data set. For human and mouse, which are considered unlabeled, we then clustered each of them to obtain unbiased reference labels with high granularity that will be mapped to the query data set.
3. Then, for each reference–query pair, we used Seurat’s FindTransferAnchors (normalization.method = “SCT”) followed by TransferData to project reference identities onto the query. Predicted labels were filtered by GetTransferPredictions with a score threshold of 0.6; cells not meeting this threshold were considered “Unassigned.”
4. Mapping success for a given direction (reference → query) was quantified as the proportion of assigned cells in the query (number assigned / total query cells). These proportions were assembled into a reference-by-query matrix (diagonal set to 1.0) and visualized as a heatmap for off-diagonal mapping rates.

We use the same label transfer strategy to map other neural identities such as coarse neural groups (dorsal excitatory, dorsal inhibitory and ventral) across species.

### Cross-species Data Integration

To jointly embed datasets into a batch-corrected cross-species molecular space, we applied Seurat’s integration framework with prior SCTransform normalization using two methods. Canonical correlation analysis (CCA) was used as the primary anchor-finding method, while reciprocal PCA (RPCA) was used as an independent validation. We chose these methods because they have been shown to perform well across distant species, achieving both strong species mixing and biology conservation while preserving cell type structure.^79^

For developmental data, we integrated cardinal-class neurons (dI1–dI6, V0–V3, MNs) across species. To balance sampling, datasets were down-sampled to the size of the dataset with the lower cell number. Each dataset was normalized with SCTransform (Seurat v5; vst.flavor = “v2”), variable features were selected (7,000 per dataset), and the anchor feature set was defined as the intersection of these features after removing ribosomal genes (Rgs* and Rpl*). We then used Seurat’s SCT-based integration workflow: PrepSCTIntegration → FindIntegrationAnchors (normalization.method = “SCT”, reduction either “cca” or “rpca”) → IntegrateData (normalization.method = “SCT”). Integrated objects were embedded with PCA (50 PCs), UMAP computed on the top PCs, and SNN graphs used for clustering.

For adult cross-species integration, each dataset was SCTransformed with appropriate technical covariates (library size and gene count; dataset-specific batch or sample terms where available). We selected 10,000 variable features per dataset, defined the anchor feature set as their intersection, and performed integration with CCA and RPCA anchors. For CCA integrations we used a higher latent dimensionality (integration and downstream on PCs 1–150); for RPCA we used PCs 1–50. The integrated objects were then embedded with PCA/UMAP and clustered for downstream visualization/analyses.

### Cross-species Similarity Index

To quantify correspondence between neural classes across species, we computed a graph-based similarity on the integrated k-nearest neighbor (kNN) graph (integrated_nn).

For a given species pair (A,B) and neural class pair (i,j), we first identified cells of class i in species A and class j in species B. For each A-cell in class i, we counted how many of its k nearest neighbors in the integrated space were B-cells of class j, divided by k to obtain a per-cell neighbor fraction. We averaged these fractions over all A-cells in class i. We repeated the procedure in the reverse direction (B→A) and reported the symmetrized similarity as the mean of the two directional averages.

This procedure yields a similarity index value in [0,1] that can be interpreted as the average fraction of a cell’s nearest neighbors that belong to the corresponding class in the other species; because most neighbors remain within-species, conserved correspondences typically score ∼0.15–0.25, whereas non-corresponding pairs are closer to zero. Because similarity indices are calculated after integration, the range of values may vary depending on the approach, making the overall trend the key criterion for evaluation. We used the same neural class labels for both species. K (size of the neighborhood) was specified when constructing a kNN graph on the integrated data with *FindNeighbors* function using *k.param* argument.

As a final note, the similarity index can be understood as an average composed of two terms. The first one reveals integration efficiency and the second one represents class mapping. Mathematically, We denote:

1. for class i in species A with *A_i_* and class j in species B, with *B_j_*.
2. A cell in*A_i_* with a, and the count of all the cells in *A_i_* with N*^i^_A_*.
3. a nearest neighbor graph with k neighbors, kNN.

Then, the similarity index is computed as:

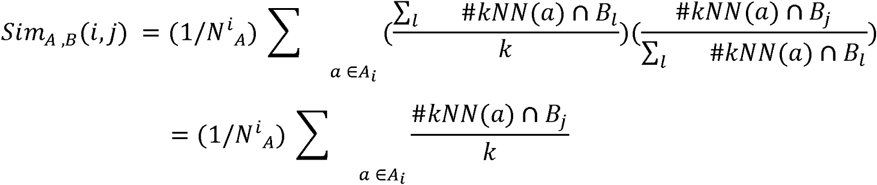

The final calculation displayed in the paper is symmetrized:

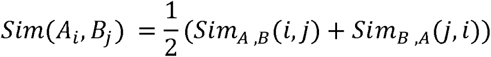

### Species Mixing Analysis

To quantify and visualize species representation within each cluster, we counted cells from frog, mouse, and human and normalized these counts by the total number of cells from each species across the dataset, thereby correcting for unequal sampling size. These normalized values were then converted into within-cluster proportions, ensuring that the contributions of frog, mouse, and human summed to one per cluster.

To enhance interpretability, proportions were square-root transformed and re-normalized, which accentuates minority contributions without altering the relative ranking of species within a cluster. Each species was assigned to one color channel of the RGB space (Human-red, Frog-green, Mouse-blue). For a given cluster, the adjusted species proportions determined the intensity of the red, green, and blue channels, producing a composite color that directly encodes the species mixture. To highlight conserved clusters with high contributions from all species, we blended such clusters toward a neutral gray (RGB value 0.75 per channel) when all three species contributed at least 5-20%. This ensured that conserved clusters appeared more muted, while species-specific or two-species-dominated clusters retained vivid colors.

The resulting colors were assigned to all cells within each cluster and projected onto the UMAP embedding for visualization. A ternary diagram using the same RGB mapping was included as a legend, with vertices corresponding to single-species clusters, edges to two-species mixtures, and the center to balanced three-species mixtures.

For the integration with zebrafish data, we collapsed mouse and human into a single “mammalian” group and reassigned color channels accordingly: zebrafish-red, frog-green, and mammals (mouse+human)-blue. The same normalization, √-transform, and gray-blending procedure was applied. This allowed direct visualization of conserved and divergent clusters across anamniotes (frog, zebrafish) and mammals within a single ternary map.

All transformations (normalization, √-transform, RGB mapping, and gray blending) were implemented in R. UMAP visualizations were generated with Seurat and ggplot2, and the ternary legend was drawn using ggtern and arranged with patchwork.

### Pearson Correlation

To quantify transcriptional similarity across species, we computed Pearson correlation coefficients on average expression profiles of neural types. For each dataset, the intersecting set of homologous genes was identified, and expression values were scaled gene-wise z-scoring them (*ScaleData* function). Cells were subset by their neural type identity, and mean expression vectors were calculated for each neural type using either the full homologous gene set or functional subsets (transcription factors, cell adhesion, ion transport, signaling). Correlation matrices were then generated between neural types of each species pair and visualized as heatmaps.

### Purity Analysis

To evaluate the contribution of different gene categories to cross-species segregation of cardinal classes, we performed clustering analyses using gene subsets. Gene lists for transcription factors, cell adhesion molecules, ion transport, and signaling receptors were obtained by mapping human homologs to Gene Ontology terms via Ensembl BioMart. For each category, variable genes from each of the lists were used as features for PCA, followed by graph-based clustering (Seurat v5, resolution = 7). Cluster purity was quantified as the proportion of cells from the most frequent cardinal class within each cluster. Distributions of purity values across all cells for each gene category were then visualized and these were then compared across gene categories and the full set of variable genes.

### Spatial Transcriptomics - Tissue Collection

The caudal half of wild-type adult *Xenopus laevis* and the dorsal portion of NF55 tadpoles — from a level just caudal to the eyes to ∼5 mm posterior to the hindlimbs — were flash frozen by immersion in a dry ice–isopentane bath for 20 seconds and stored at –80 °C until further processing. Prior to sectioning, tissues were embedded in optimal cutting temperature (OCT) compound (manufacturer) within molds placed on dry ice. Cryosections (10 µm) were collected onto glass slides supplied by Resolve Biosciences (Resolve BioSciences GmbH, Monheim am Rhein, Germany). Samples included four NF55 tadpoles, with sections collected at brachial, thoracic, and lumbar spinal levels, and three adult males, from which only lumbar spinal cord sections were obtained. Slides were stored at –80 °C for less than one week before shipment on dry ice to Resolve Biosciences for downstream processing. Two probe sets (developmental and adult) were designed using the Resolve Biosciences algorithm (**Table S1**; **Table S2**). Overall, 85–90% of sections passed quality control, while the remainder were excluded due to damaged anatomy or suboptimal probe detection.

### Spatial Transcriptomics – Imaging

Spatial transcriptomic imaging was performed using the Resolve Biosciences Molecular Cartography™ platform on a Zeiss Celldiscoverer 7 equipped with a 50× Plan-Apochromat water-immersion objective (NA 1.2) and a 0.5× magnification changer, yielding a final 25× magnification. Images were acquired using standard CD7 LED illumination, appropriate filter sets, and a Zeiss Axiocam Mono 712 CMOS camera (3.45 µm pixel size). Each region was imaged across eight hybridization rounds, with two fluorescence channels per round, producing 16 image stacks per position. Imaging was fully automated via a Python-based control script using the Zeiss ZEN API.

### Spatial Transcriptomics - Spot Segmentation and Preprocessing

Raw images were processed and decoded using Resolve’s proprietary Molecular Cartography software. Background correction, signal detection, and 3D spot segmentation were carried out following Resolve’s standard pipeline, which utilizes algorithms implemented in Java (based on the ImageJ framework) and C++ (using the libpointmatcher library for point-cloud alignment). Detected fluorescence maxima across z-stacks were grouped into 3D features, filtered by brightness and local background intensity, and used to generate point clouds representing individual transcript candidates. Point clouds from each imaging round were aligned to a reference round using iterative closest-point registration to achieve sub-pixel accuracy. Decoded signals were then assigned to genes according to hybridization code profiles, and quality metrics were applied to remove low-confidence or non-specific detections. The resulting transcript coordinates were used for downstream spatial analysis.

### Spatial Transcriptomics - Cell Segmentation

Count tables of molecules containing x and y coordinates were obtained from Resolve. We used Baysor (Petukhov et al Nat Biotech 2021, https://www.nature.com/articles/s41587-021-01044-w) version 0.5.2 to segment cells using the spatial transcriptomics data alone. Briefly, Baysor models a cell as an 2D ellipsoid (µ, Σ) and a vector of gene expression frequencies *f*. The likelihood of generating one molecule *i* with gene *g_i_* at location xi from a cell *z_i_* is: *Pr(g_i_, x_i_); N(x_i_|µ_Zi_, Σ_Zi_) Mul(1|f)*.

To capture the spatial clustering tendencies in the data (e.g., molecules that are close in space tend to belong to the same cell), Baysor uses Markov Random Field priors using the Potts model. Baysor also includes a model for the background component and prior information of expected cell size and minimal number of molecules per cell. The full probabilistic model is found in Petukhov et al 2021, and the latent variables are inferred using a Stochastic EM algorithm with variational optimization. We performed cell segmentation using the Baysor software, setting minimum number of molecules m to 10 and expected cell size s to 18:
baysor run -x x -y y -z z -g gene -m 10 -s 18 --plot -o ${outputdir} --save-polygons geojson ${inputfile}

### Spatial Transcriptomics - Label Transfer and Visualization

After cell segmentation, data was converted into count matrices with cell coordinates and slide metadata.

For adult frog spatial data, probe identifiers that did not match snRNA-seq gene symbols were remapped (e.g., chat.L→LOC108696296, meis2.L→LOC108699226, tcf4.L→Xelaev18011072m), while rxfp2 was excluded due to inconsistent signals. A Seurat object was created from the harmonized matrix and normalized with SCTransform. Coarse identities (Neurons vs non-neuronal) were first assigned by SCT-based label transfer from the integrated adult frog reference, using a prediction score threshold of 0.5. Neuronal cells were then mapped to broad classes (excitatory, inhibitory, or motor neurons) with a curated neuron reference (score ≥0.5), and subsequently resolved into finer families (e.g., Exc_Cpne4/Bnc2, Exc_Skor1, Exc_Npy, Exc_Tcf4/Penk, Exc_Vent; Inh_Vent, Inh_Ntn1, Inh_Tcf4/Nfi, Inh_Nos1, Inh_Rorb, Inh_Other) by SCT-based transfer (score ≥0.3). Assignments were validated with marker-based dot plots, and final identities were projected onto cell centroids for spatial maps and normalized density profiles along the dorsal–ventral axis.

For developmental frog spatial data, probe identifiers were harmonized to snRNA-seq gene symbols (e.g., chat.L→LOC108696296, hoxc6.L→LOC108708639, olig2.L→LOC108707878, otp.L→LOC108717966, pou6f2.L→LOC108719010, sox2.L_10.1→sox2.L). A Seurat object was generated (SCTransform; minmol = 6, scale = 30) and sections with abnormal signal or cells outside the gray matter were excluded; slides were manually rotated to align dorsal–ventral axes across animals and spinal levels. Cell identities were assigned by SCT-based label transfer from an NF54 scRNA-seq reference, yielding coarse categories (roof plate, floor plate, progenitors/differentiating neurons, neurons, oligodendrocytes, other) at a prediction score threshold of 0.5. Neurons were then subset, and assignment to cardinal classes (dI1–dI6, V0–V3, MNs) was performed using a curated panel of developmental markers (e.g., barhl1/2, lhx2/9, hmx3, isl1/2, tlx3, sall3, lmx1b, wt1, dmrt3, evx1/2, en1, foxp2, lhx3, vsx2, sim1, tal1, gata3, nkx2-2). To reduce spurious assignments, spatial cells were first filtered to retain only those expressing at least one marker gene from this panel. For each cardinal class cluster in the scRNA-seq reference, average log-expression profiles were computed across the marker set, and Pearson correlations were calculated between these reference profiles and each spatial cell. Each cell was then assigned to the class with the highest correlation, and MNs were merged back from the coarse stage to yield the final set of cardinal class identities.

### Spatial transcriptomics – Projecting Conserved and Divergent Populations

To visualize the spatial distribution of conserved versus divergent neuronal populations, we used the three-species integrated Seurat objects (frog, mouse, human) to derive conserved/divergent cluster identities. For each domain (dorsal excitatory, dorsal inhibitory, and ventral/midline), clusters were annotated as Conserved or Divergent based on their cross-species mixing patterns in the integrated dataset (**Fig. 4**). These annotations were propagated back to the original single-species datasets (frog and mouse – Russ et al. dataset) and used for spatial mapping.

For frog, conserved/divergent labels from the integrated object were transferred to the original single-cell neuronal subset and projected onto Molecular Cartography spatial data using Seurat label transfer (FindTransferAnchors, normalization.method = “SCT”; TransferData, prediction.assay = TRUE). Prediction scores for Divergent neurons were plotted across spatial coordinates of the Xenopus adult spinal cord sections (**Fig. 4J**).

For mouse, Visium data from Kathe et al. (GEO: GSE184369) were first processed to identify neuronal spots using label transfer from the Russ et al. scRNA-seq dataset. The resulting neuronal subset was then used as a query for projecting conserved/divergent identities from the Russ et al. mouse dataset via FindTransferAnchors (normalization.method = “LogNormalize”) and TransferData. Divergent prediction scores were visualized as spatial gradients over Visium coordinates to highlight laminar localization of divergent neurons (**Fig. 4J**).

## Data and Materials Availability

Upon publication, all data will be made available in the main text or the supplementary materials and sequencing data will be deposited in GEO and available for download.

## Code Availability

All code used for the analysis of single-cell and spatial transcriptomic data will be made available on GitHub (https://github.com/yuriignatyev) upon publication.

## Acknowledgements

We would like to thank the members of the Sweeney Lab for discussion and support; Andrey Bydanov for technical assistance with single-cell sequencing processing; and Jay Bikoff, Nikos Konstantinides, Maria Tosches, and Graziana Gatto for comments on the manuscript.

## Author Contributions

L.B.S. led and coordinated the project. Y.I., M.I.G. and L.B.S. designed the study. Y.I. led all computational analyses of single cell and spatial transcriptomic data, supported by guidance of M.I.G. and L.B.S., and with assistance from M.S., J.Y., and L.P. Y.I., M.S. and L.B.S collected scRNAseq and snRNAseq data. S.P., in collaboration with Resolve and Y.I., performed all spatial transcriptomic experiments in tadpoles and adult frogs. T-Y.L. provided initial, unpublished methodologies for dissociating amphibian single cells, from the laboratory of E.M.T. Y.I., M.I.G. and L.B.S. wrote the manuscript. S.P. and A.J.L. edited the manuscript and provided critical feedback on the project.

## Supplementary Materials

Figs. S1 to S20.

Tables S1 to S4.

