## Supplementary material for "Innovations in spinal cord cell type heterogeneity across vertebrate evolution": Figs. S1-S20

[1]IST Austria, Klosterneuburg, Austria; [2]Heidelberg University, Heidelberg, Germany; [3]Research Institute of Molecular Pathology (IMP), Vienna BioCenter (VBC), Vienna, Austria; [4]Harvard Medical School, Department of Systems Biology, Boston, MA; [5]Spinal Circuits and Plasticity Unit, National Institute of Neurological Disorders and Stroke, Bethesda, MD; [6]Allen Institute for Brain Science, Seattle, WA; [7] University of Washington, Department of Statistics, Seattle, WA

*corresponding author

#### Supplementary Figures

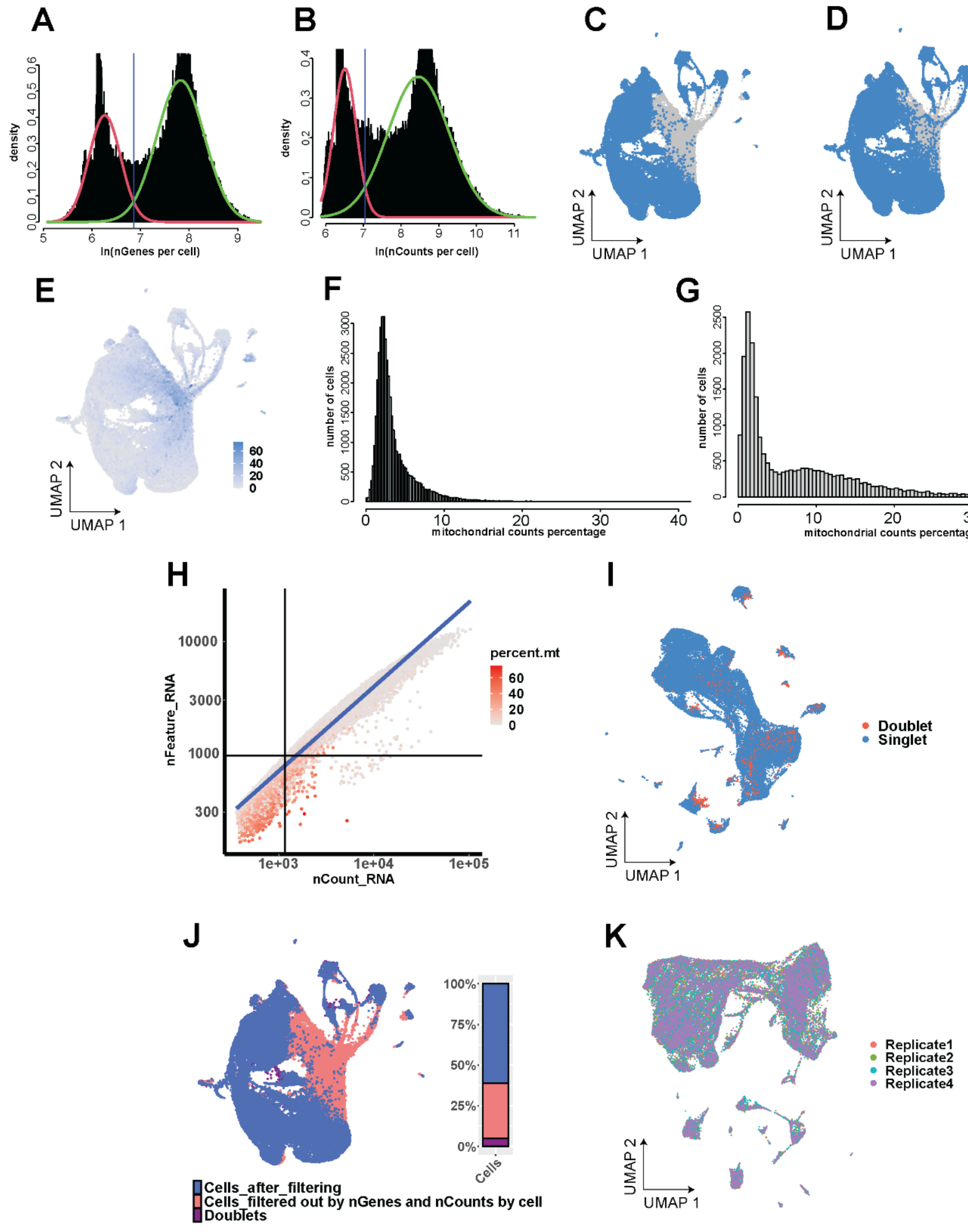


**Figure S1**

**Fig. S1. Quality control metrics for the amphibian spinal cord single-cell atlas at NF stage 54.**

**A–B.** Bimodal distributions of log-normalized gene numbers (**A**) and counts (**B**) per cell. The blue line indicates the threshold used to distinguish high- from low-quality cells.

**C–E.** UMAP representation of all cells in the dataset. Cells passing the threshold for number of genes (**C**) and counts (**D**), corresponding to the distributions in (**A**) and (**B)**, are shown in blue. In **E**, cells are color-coded by the percentage of mitochondrial reads per cell (light blue). Cells with a high proportion of mitochondrial reads overlap with the low-quality cells filtered out in (**C**) and (**D**).

**F–G.** Distribution of mitochondrial read percentages in cells that passed quality thresholds (**F**) and in filtered-out cells (**G**).

**H.** Scatter plot of the number of reads (nCount_RNA) versus the number of detected genes (nFeature_RNA) per cell, colored by mitochondrial read percentage. Lines indicate the thresholds applied to filter cells based on gene and read counts. Cells failing these thresholds largely correspond to those with elevated mitochondrial read percentages.

**I.** UMAP representation of cells passing QC filters, with doublets (red) and non-doublets (blue) identified using *DoubletFinder*.

**J.** UMAP representation of all cells in the dataset. Cells removed due to low gene or count thresholds are shown in red, doublets in purple, and high-quality cells that passed all filtering steps in blue.

**K.** UMAP representation of cells after all QC filtering steps, color-coded by technical replicates, shows good mixing of samples within the UMAP space.

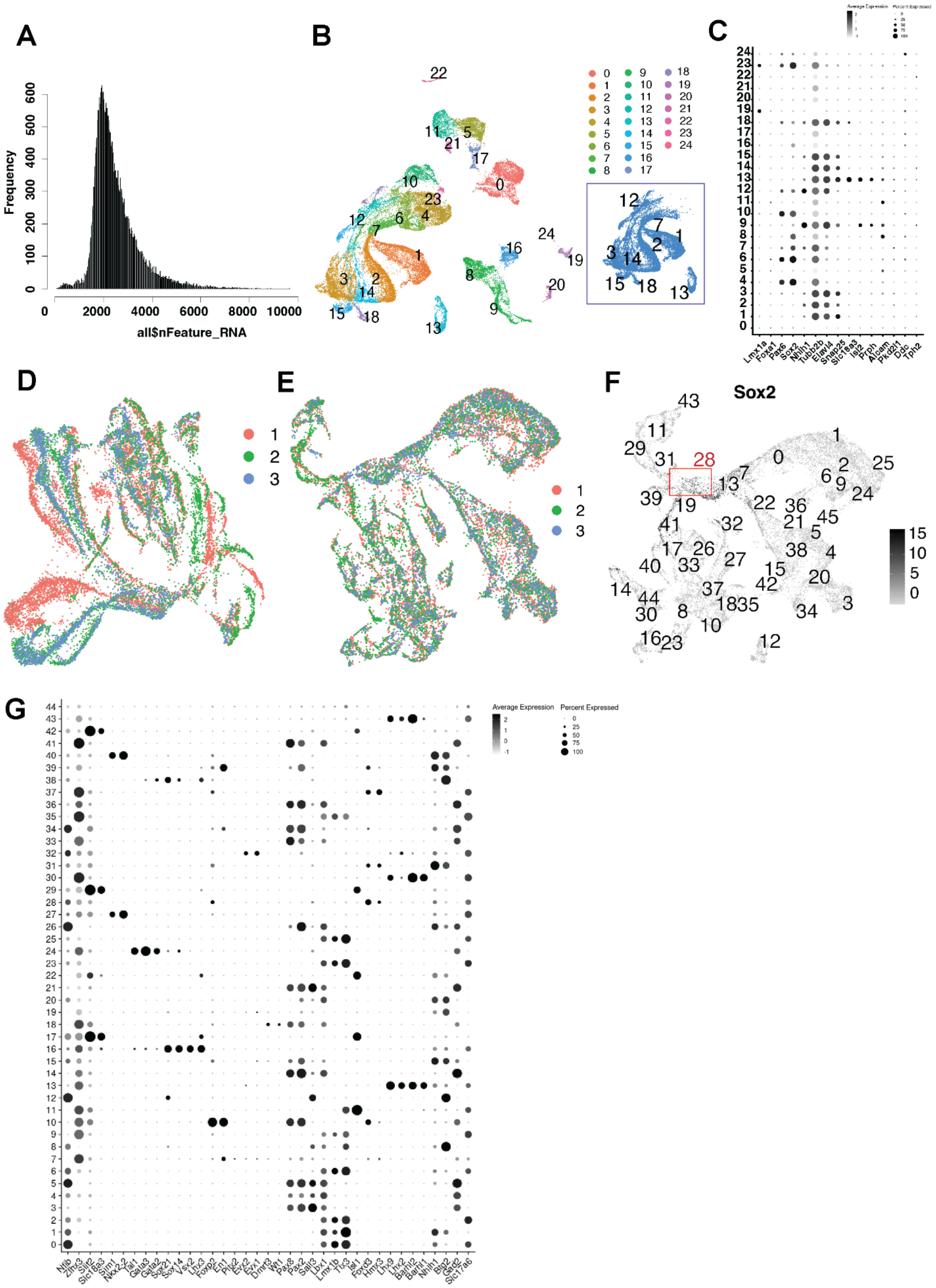
A
B
C
D
E
F
G
Frequency
all$nFeature_RNA
Sox2
Average Expression
Percent Expressed
Lmx1a
Foxa1
Pax6
Sox2
Nhlh1
Tubb2b
Elavl4
Snap25
Slc18a3
Isl2
Prph
Alcam
Ddc
Tph2

**Figure S2**

**Fig. S2. Analysis of mouse embryonic spinal cord neural data from *Delile et al. 2019.***

**A.** Distribution of genes per cell in *Delile et al. 2019* mouse embryonic data set.

**B.** UMAP of QC-filtered cellular transcriptomes, color-coded by unbiased clusters. Inset highlights clusters identified as neurons (blue).

**C.** Dotplot showing neural and neural tissue-related marker gene expression for clusters shown in **B**.

**D-E.** UMAP representation of neural expression profiles labelled by replicate before (**D**) and after batch correction (**E**).

**F**. Same data as in **E** colored by expression of Sox2 and numbered according to unbiased clusters. Cluster 28 exhibits expression of Sox2 indicating progenitor identity.

**G.** Dot plot showing expression of cardinal class markers across the unbiased clusters from **F**, after removal of cluster 28, which consisted of progenitor cells.

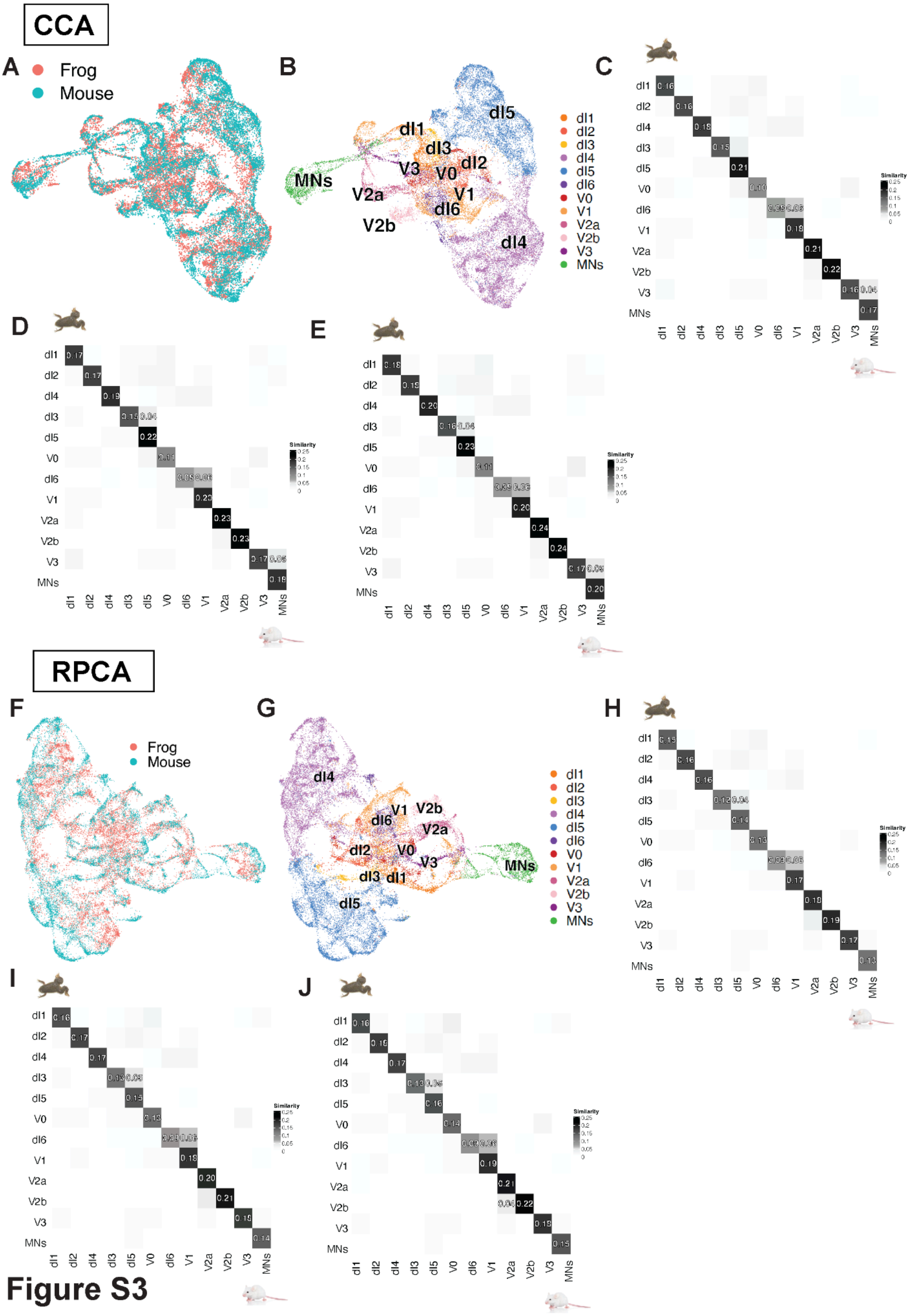
CCA
A
Frog
Mouse
B
dI1
dI2
dI3
dI4
dI5
dI6
V0
V1
V2a
V2b
V3
MNs
C
D
E
Similarity
RPCA
F
G
H
I
J

**Figure S3**

**Fig. S3. Integration of frog and mouse developmental data as in Fig. 2E using multiple methods.**

**A-B.** UMAP-representation of frog and mouse integrated neural data using canonical correlation analysis (CCA) after performing SCTransform-based normalization, labelled by species (**A**) and cardinal classes (**B**).

**C-E.** Heatmap representing analysis of molecular convergence across neural types in frog (vertical) and mouse (horizontal) based on the proportion of overlapping mutual nearest neighbours by species across neural types with size of neighbourhood 15 (**C**), 20 (**D**) and 25 (**E**) in the KNN-graph of the integrated with CCA cross-species data.

**F-G.** UMAP-representation of frog and mouse integrated neural data using reciprocal PCA (RPCA) after performing SCTransform-based normalization, labelled by species (**F**) and cardinal classes (**G**).

**H-J.** Heatmap representing analysis of molecular convergence across neural types in frog (vertical) and mouse (horizontal) based on the proportion of overlapping mutual nearest neighbours by species across neural types with size of neighbourhood 15 (**H**), 20 (**I**) and 25 (**J**) in the KNN-graph of the integrated with RPCA cross-species data.

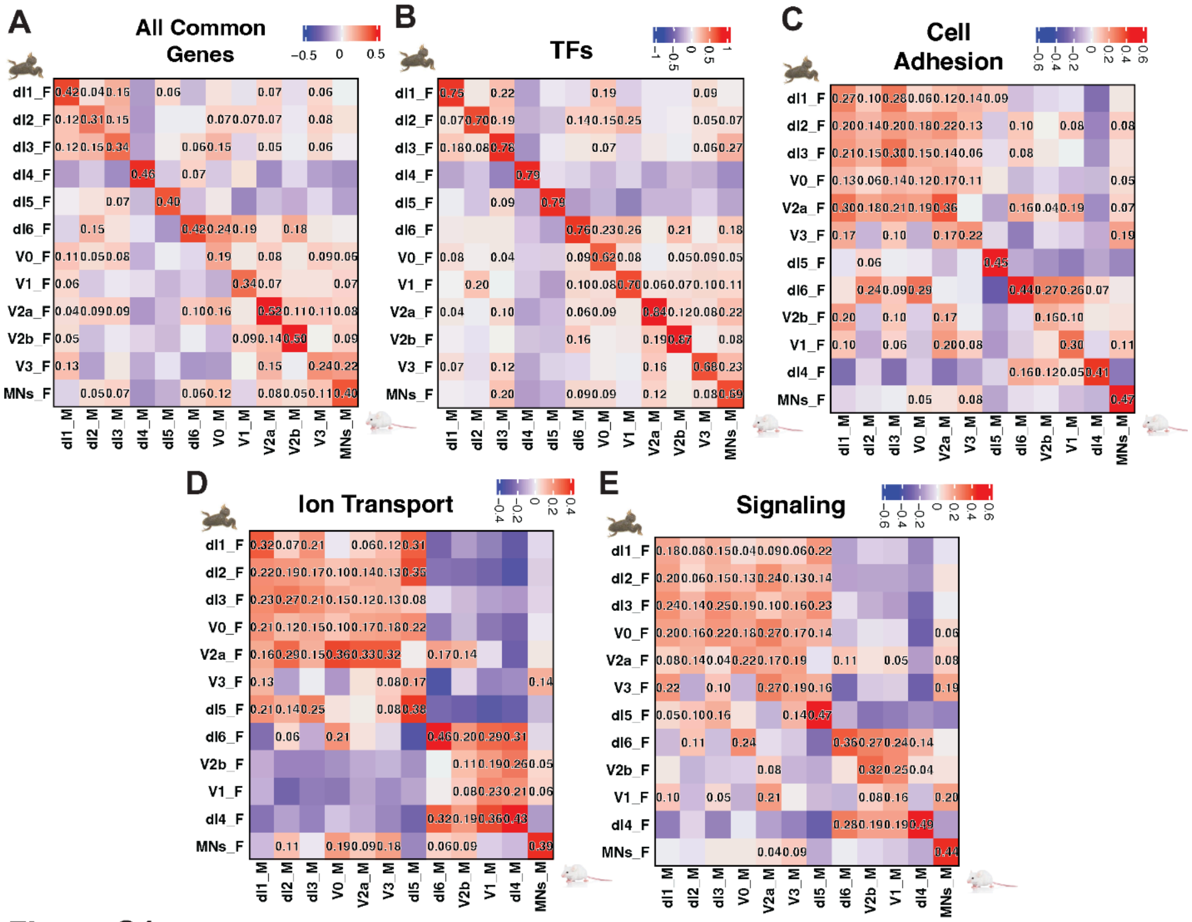
A
All Common Genes
B
TFs
C
Cell Adhesion
D
Ion Transport
E
Signaling

**Figure S4**

**Fig. S4. Pearson correlation of gene expression profiles across neural types between developing spinal cord of frog and mouse arranged by gene categories.**

**A–E.** Heatmaps showing Pearson correlation coefficients between frog and mouse neural type expression profiles, calculated using different gene sets: all genes (13,281 genes; **A**), transcription factors (948 genes; **B**), cell adhesion genes (758 genes; **C**), ion transport genes (646 genes; **D**), and signaling genes (614 genes; **E**).

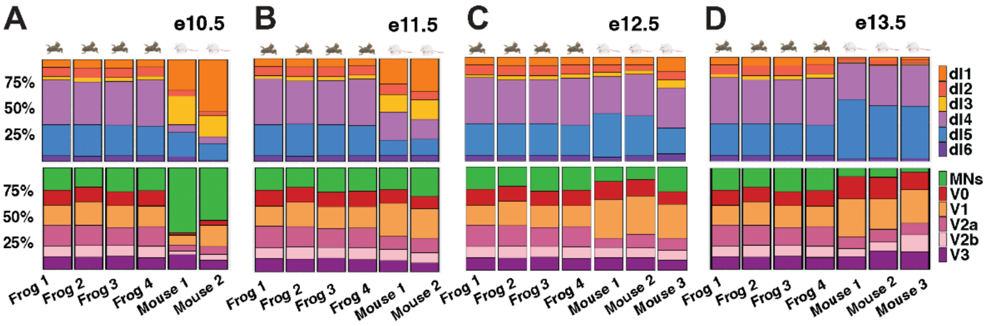
A
e10.5
B
e11.5
C
e12.5
D
e13.5
75%
50%
25%
dI1
dI2
dI3
dI4
dI5
dI6
MNs
V0
V1
V2a
V2b
V3
Frog 1
Frog 2
Frog 3
Frog 4
Mouse 1
Mouse 2
Mouse 3

### Figure S5

**Fig. S5. Spinal cord cell type proportions in frog (stage NF54) and mouse (stages E10.5–E13.5).**

**A–D.** Proportions of dorsal (top) and ventral (bottom) cardinal classes in the developing frog spinal cord at stage NF54 and in mouse spinal cords at stages E10.5 (**A**), E11.5 (**B**), E12.5 (**C**), and E13.5 (**D**), respectively.

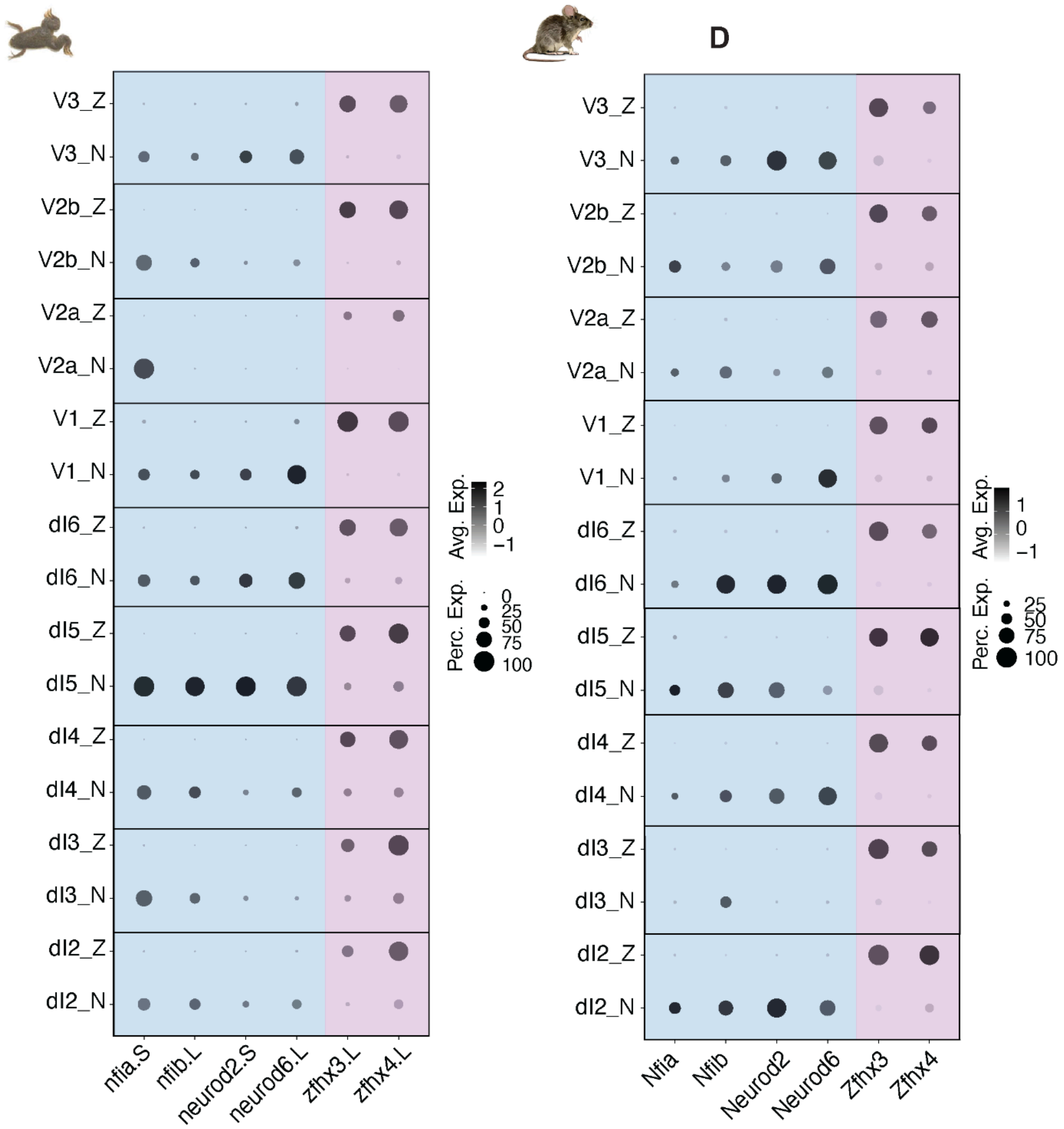
D
V3_Z
V3_N
V2b_Z
V2b_N
V2a_Z
V2a_N
V1_Z
V1_N
dI6_Z
dI6_N
dI5_Z
dI5_N
dI4_Z
dI4_N
dI3_Z
dI3_N
dI2_Z
dI2_N
nfia.S
nfib.L
neurod2.S
neurod6.L
zfhx3.L
zfhx4.L
Avg. Exp.
2
1
0
−1
Perc. Exp.
0
25
50
75
100
Nfia
Nfib
Neurod2
Neurod6
Zfhx3
Zfhx4
Avg. Exp.
1
0
−1
Perc. Exp.
25
50
75
100

**Figure S6**

**Fig. S6. Division of cardinal class subtypes by temporal *N*- and *Z*-genes associated with neuronal birth order in developing frog and mouse spinal cords.**
Dot plots show the expression of genes linked to early/mid birth order (*Z*-genes: *zfhx3, zfhx4*; red) and late birth order (*N*-genes: *nfia, nfib, neurod2, neurod6*; purple) in subpopulations of frog NF54 (left) and mouse E12.5 (right) cardinal classes.

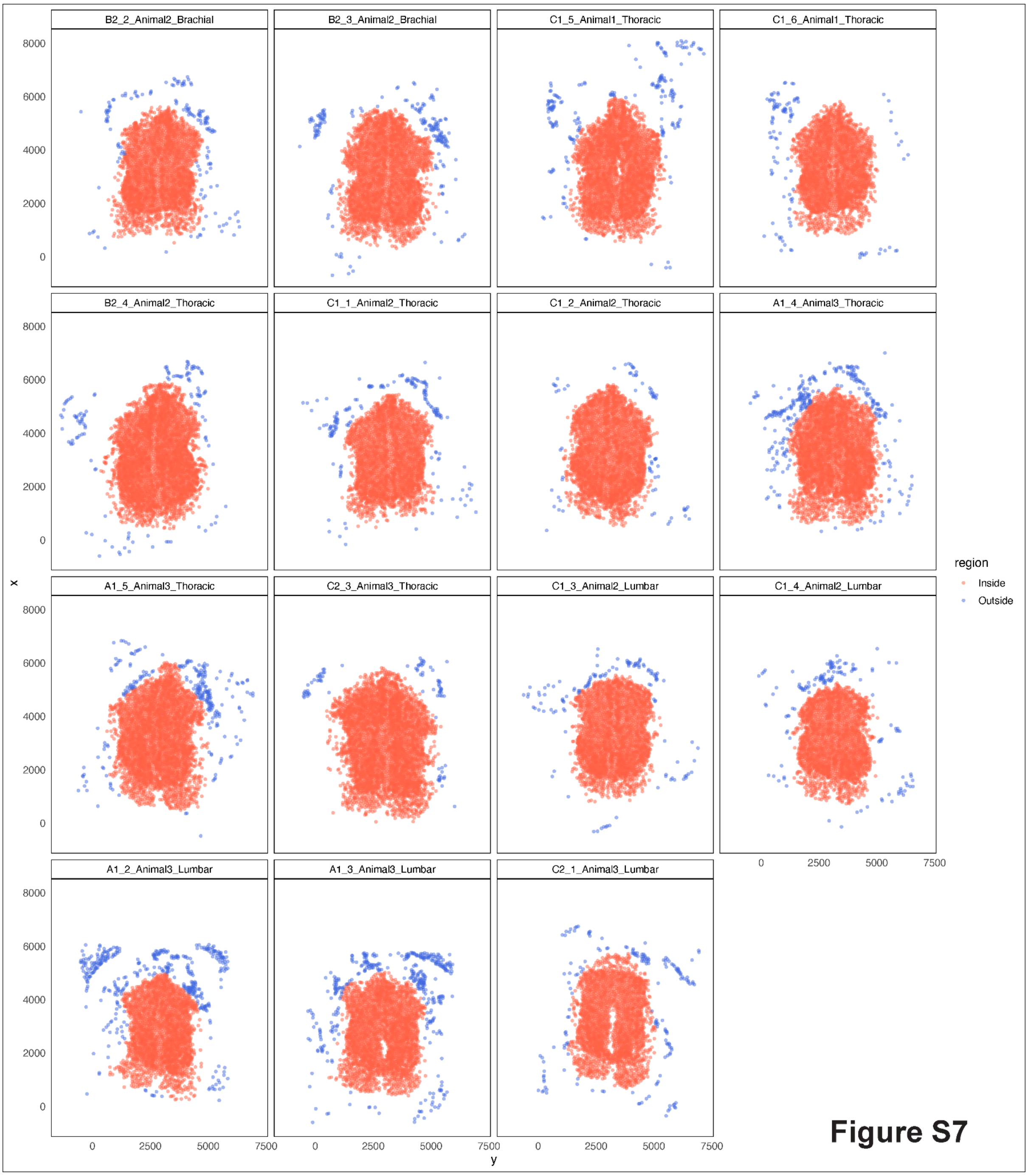
B2_2_Animal2_Brachial
B2_3_Animal2_Brachial
C1_5_Animal1_Thoracic
C1_6_Animal1_Thoracic
B2_4_Animal2_Thoracic
C1_1_Animal2_Thoracic
C1_2_Animal2_Thoracic
A1_4_Animal3_Thoracic
A1_5_Animal3_Thoracic
C2_3_Animal3_Thoracic
C1_3_Animal2_Lumbar
C1_4_Animal2_Lumbar
A1_2_Animal3_Lumbar
A1_3_Animal3_Lumbar
C2_1_Animal3_Lumbar
x
y
0
2000
4000
6000
8000
0
2500
5000
7500
region
Inside
Outside

**Figure S7**

**Fig. S7. Frog spinal cord sections with segmented cells, highlighting cells within the gray matter used for further analysis (red) and excluded cells (blue).**

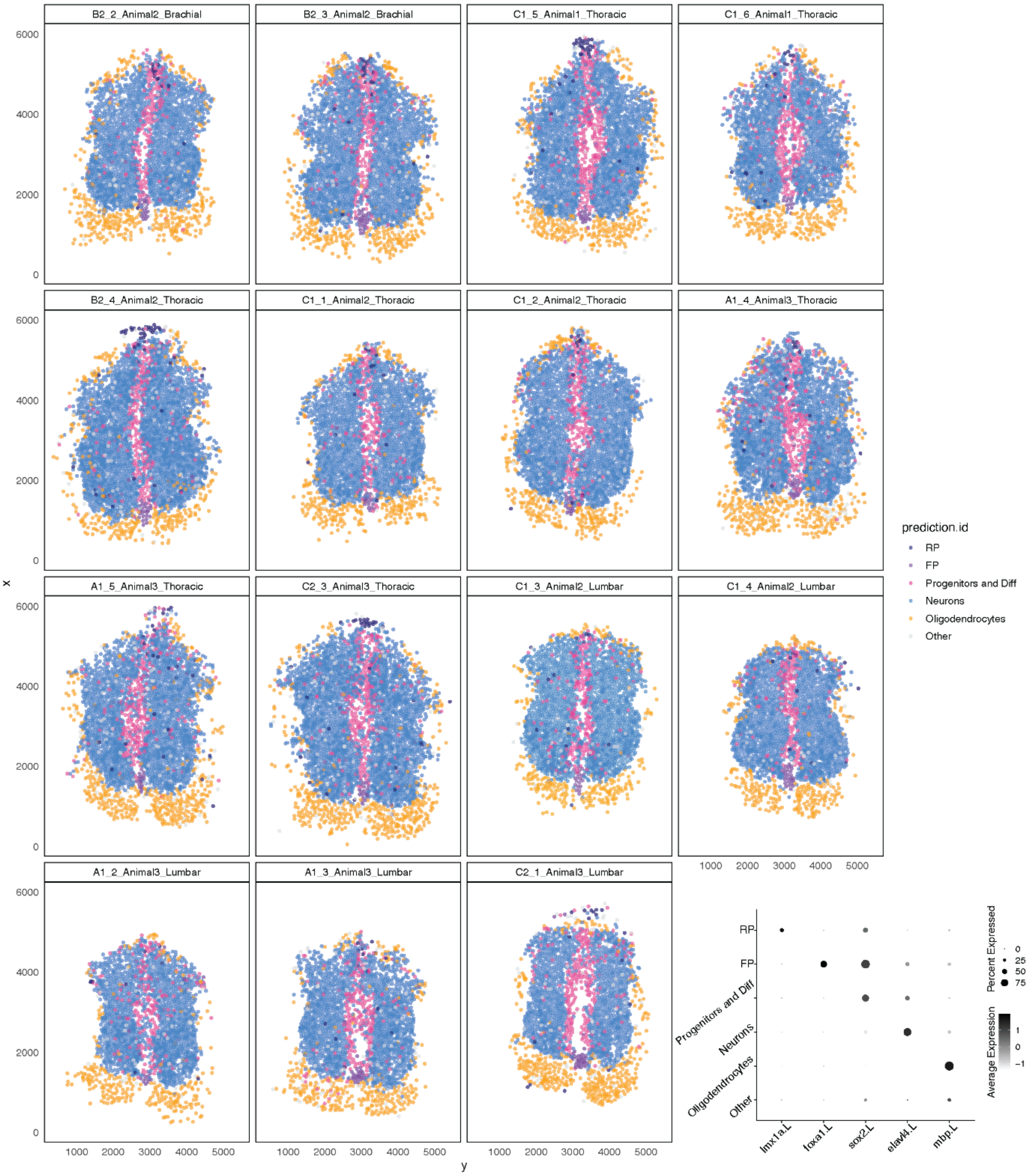

B2_2_Animal2_Brachial
B2_3_Animal2_Brachial
C1_5_Animal1_Thoracic
C1_6_Animal1_Thoracic
B2_4_Animal2_Thoracic
C1_1_Animal2_Thoracic
C1_2_Animal2_Thoracic
A1_4_Animal3_Thoracic
A1_5_Animal3_Thoracic
C2_3_Animal3_Thoracic
C1_3_Animal2_Lumbar
C1_4_Animal2_Lumbar
A1_2_Animal3_Lumbar
A1_3_Animal3_Lumbar
C2_1_Animal3_Lumbar
x
y
0
2000
4000
6000
1000
2000
3000
4000
5000
prediction.id
RP
FP
Progenitors and Diff
Neurons
Oligodendrocytes
Other
lmx1a.L
foxa1.L
sox2.L
elavl4.L
mbp.L
Percent Expressed
0
25
50
75
Average Expression
1
0
−1


### Figure S8

**Fig. S8. Coarse cell types in the frog spinal cord spatial dataset.**

**A.** Frog spinal cord sections with segmented cells labeled by coarse cell type identity via label transfer using the single-cell data as reference.

**B.** Dot plot showing the expression of coarse cell type markers represented in the spatial dataset, across segmented and labeled cells from all frog spinal cord sections.

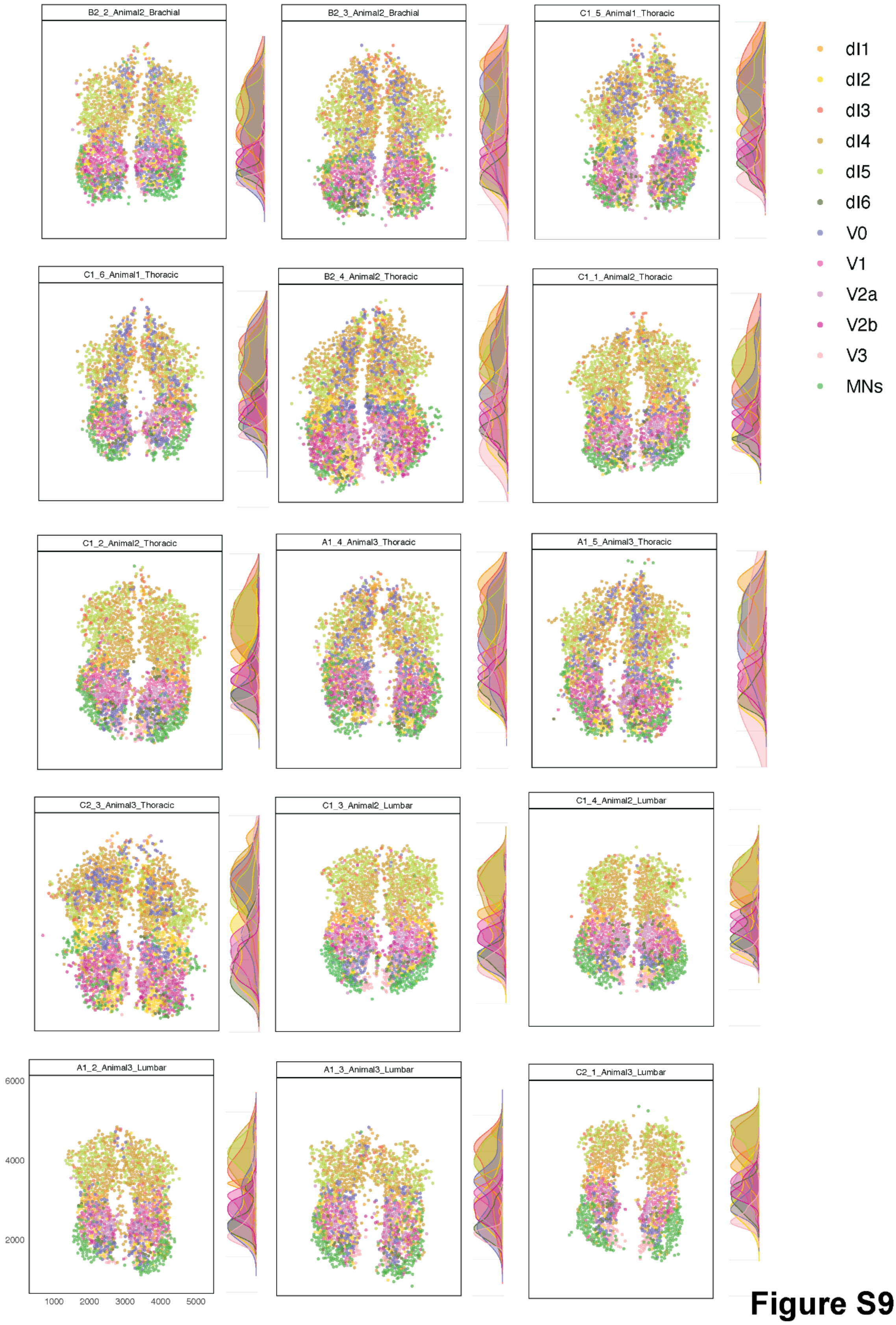
B2_2_Animal2_Brachial
B2_3_Animal2_Brachial
C1_5_Animal1_Thoracic
C1_6_Animal1_Thoracic
B2_4_Animal2_Thoracic
C1_1_Animal2_Thoracic
C1_2_Animal2_Thoracic
A1_4_Animal3_Thoracic
A1_5_Animal3_Thoracic
C2_3_Animal3_Thoracic
C1_3_Animal2_Lumbar
C1_4_Animal2_Lumbar
A1_2_Animal3_Lumbar
A1_3_Animal3_Lumbar
C2_1_Animal3_Lumbar
6000
4000
2000
1000
2000
3000
4000
5000
dI1
dI2
dI3
dI4
dI5
dI6
V0
V1
V2a
V2b
V3
MNs

**Figure S9**

**Fig. S9. Frog spinal cord sections with segmented cells labeled by cardinal class, based on correlation with expression profiles using single-cell dataset as reference.** Corresponding density plots illustrate the scaled spatial distribution of cardinal classes along the dorso–ventral axis.

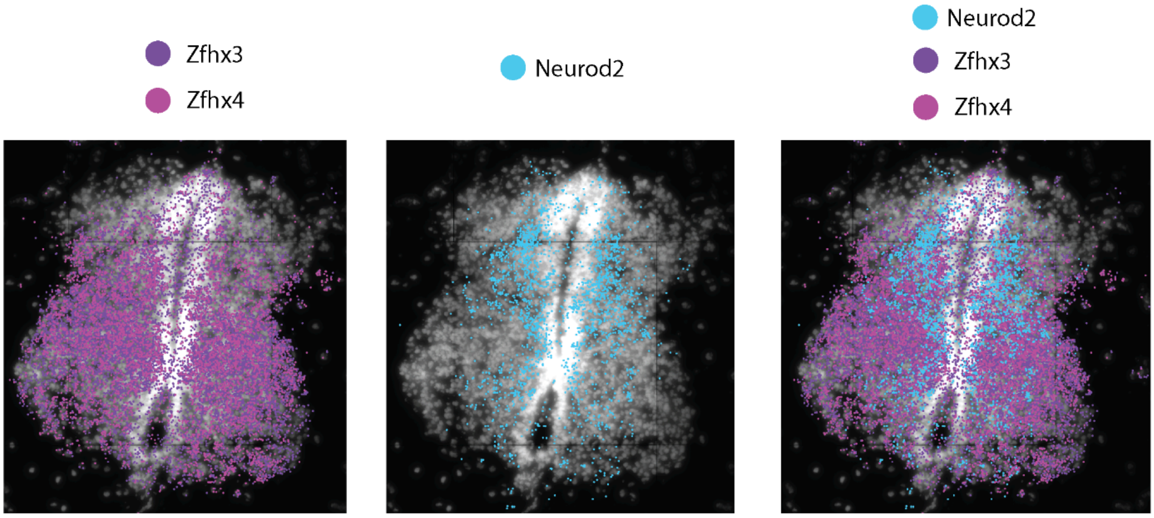


**Figure S10**

**Fig. S10. Representative spinal cord section of Neurod2 (blue) and Zfhx3/4 (purple) revealed that the N- and Z- divisions segregate cardinal classes into lateral and medial populations in developing frog, as in mice.**

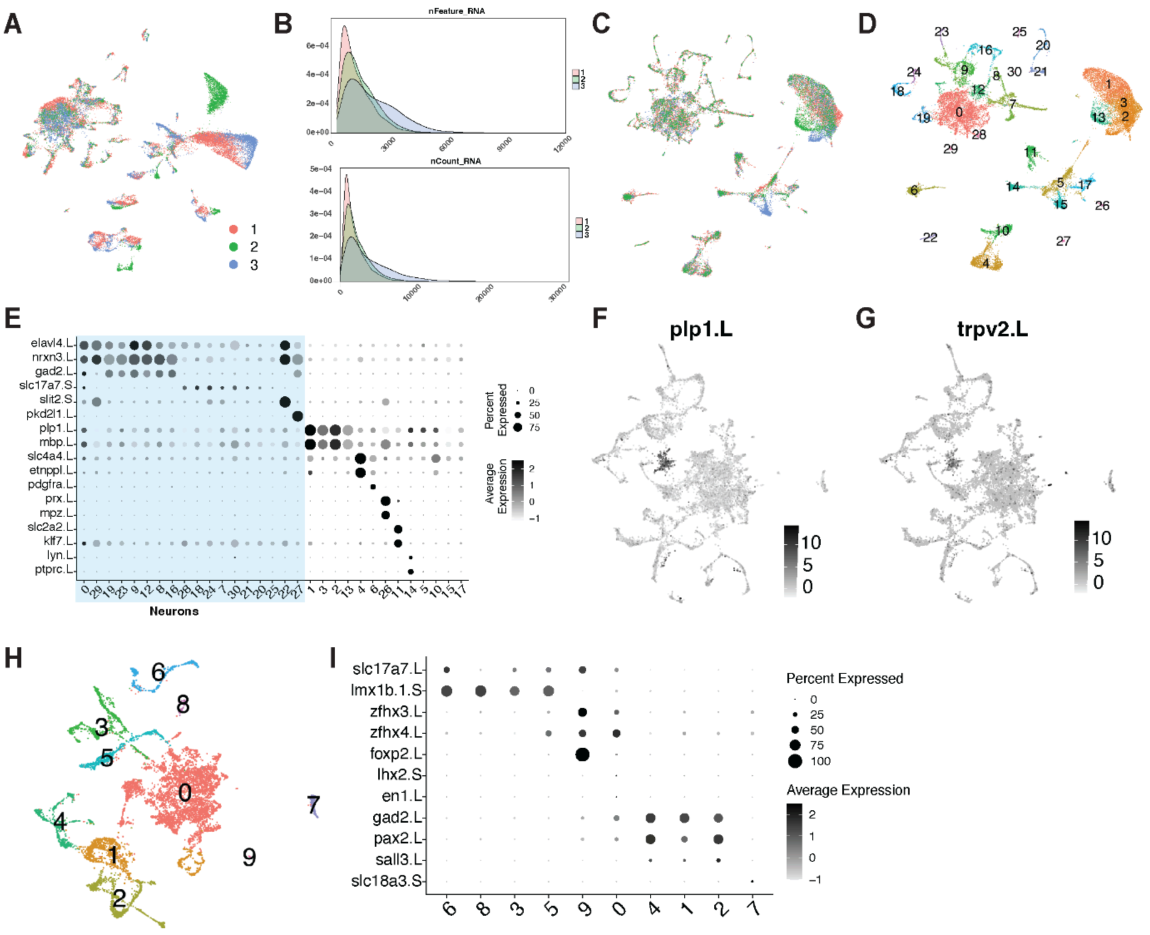
A
B
nFeature_RNA
nCount_RNA
C
D
E
elavl4.L
nrxn3.L
gad2.L
slc17a7.S
slit2.S
pkd2l1.L
plp1.L
mbp.L
slc4a4.L
etnppl.L
pdgfra.L
prx.L
mpz.L
slc2a2.L
klf7.L
lyn.L
ptprc.L
Neurons
Percent Expressed
Average Expression
F
plp1.L
G
trpv2.L
H
I
slc17a7.L
lmx1b.1.S
zfhx3.L
zfhx4.L
foxp2.L
lhx2.S
en1.L
gad2.L
pax2.L
sall3.L
slc18a3.S
Percent Expressed
Average Expression

### Figure S11

**Fig. S11. Quality control of the amphibian adult spinal cord single-nuclei atlas and neural identification.**

**A.** UMAP representation displaying single-nuclei data before integration colored by replicate.
**B.** Distributions of number of genes (top) and reads (bottom) per nuclei colored by replicate.
**C-D.** UMAP representation of single-nuclei data after integration colored by animal (**C**) and unbiased cluster (**D**).
**E.** Dot plot showing normalized expression of marker genes used to assign cell type identities across unbiased clusters. Clusters identified as neurons are highlighted in blue.
**F–G.** Feature plots showing evidence of contamination in the filtered neuronal dataset, with expression of oligodendrocyte markers (**F**) and sensory neuron markers (**G**).
**H.** UMAP representation of the filtered neuronal dataset, color-coded by unbiased clusters.
**I.** Dot plot showing marker gene expression across unbiased clusters in the filtered neuronal dataset.

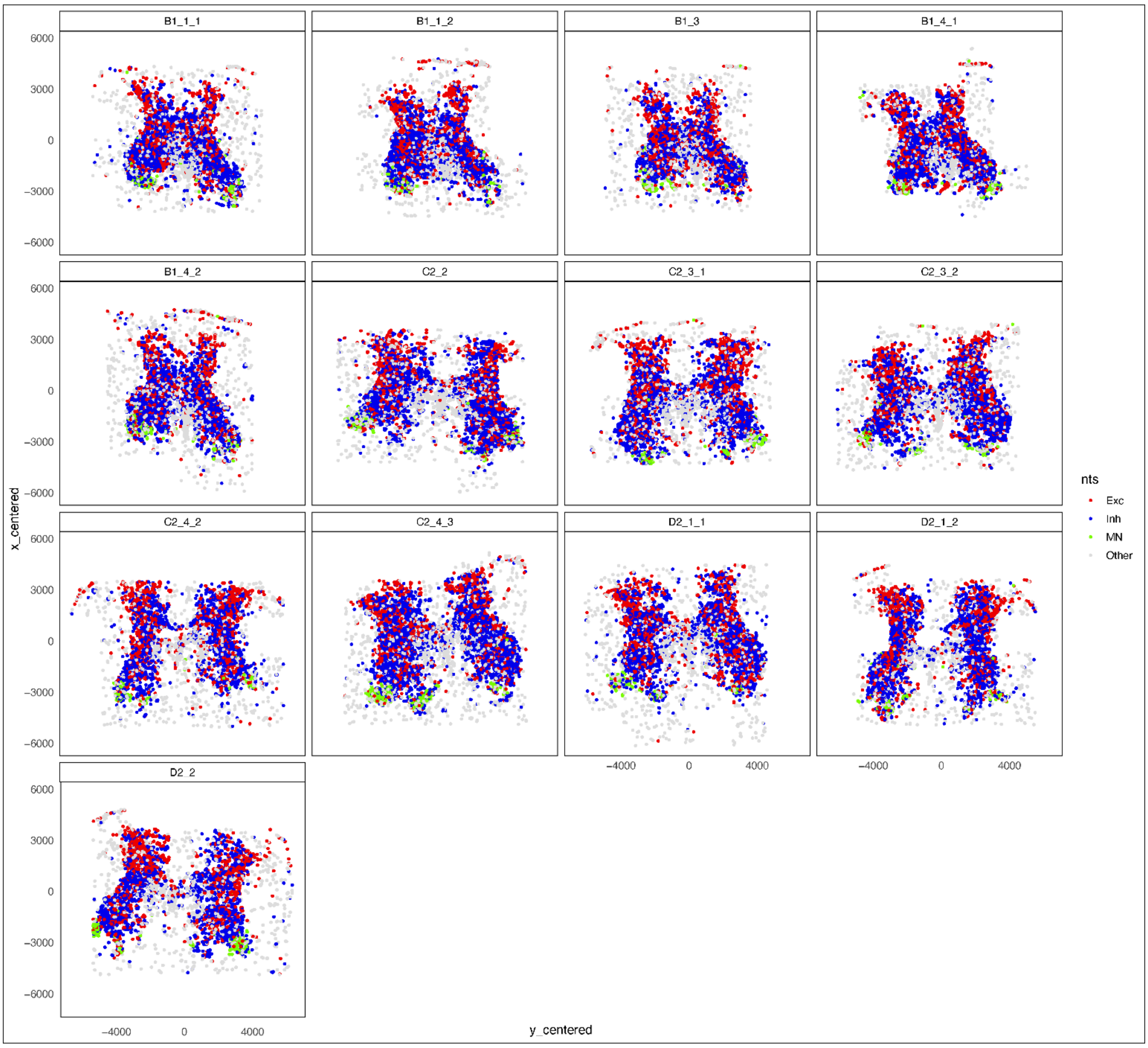
B1_1_1
B1_1_2
B1_3
B1_4_1
B1_4_2
C2_2
C2_3_1
C2_3_2
C2_4_2
C2_4_3
D2_1_1
D2_1_2
D2_2
6000
3000
0
−3000
−6000
−4000
0
4000
x_centered
y_centered
nts
Exc
Inh
MN
Other

**Figure S12**

**Fig. S12. Adult frog spinal cord sections showing segmented inhibitory (blue), excitatory (red), and motor (green) neurons.** Neuronal identities in the spatial dataset were assigned by label transfer from the single-nucleus reference dataset. Cells not labeled as neurons are shown in gray.

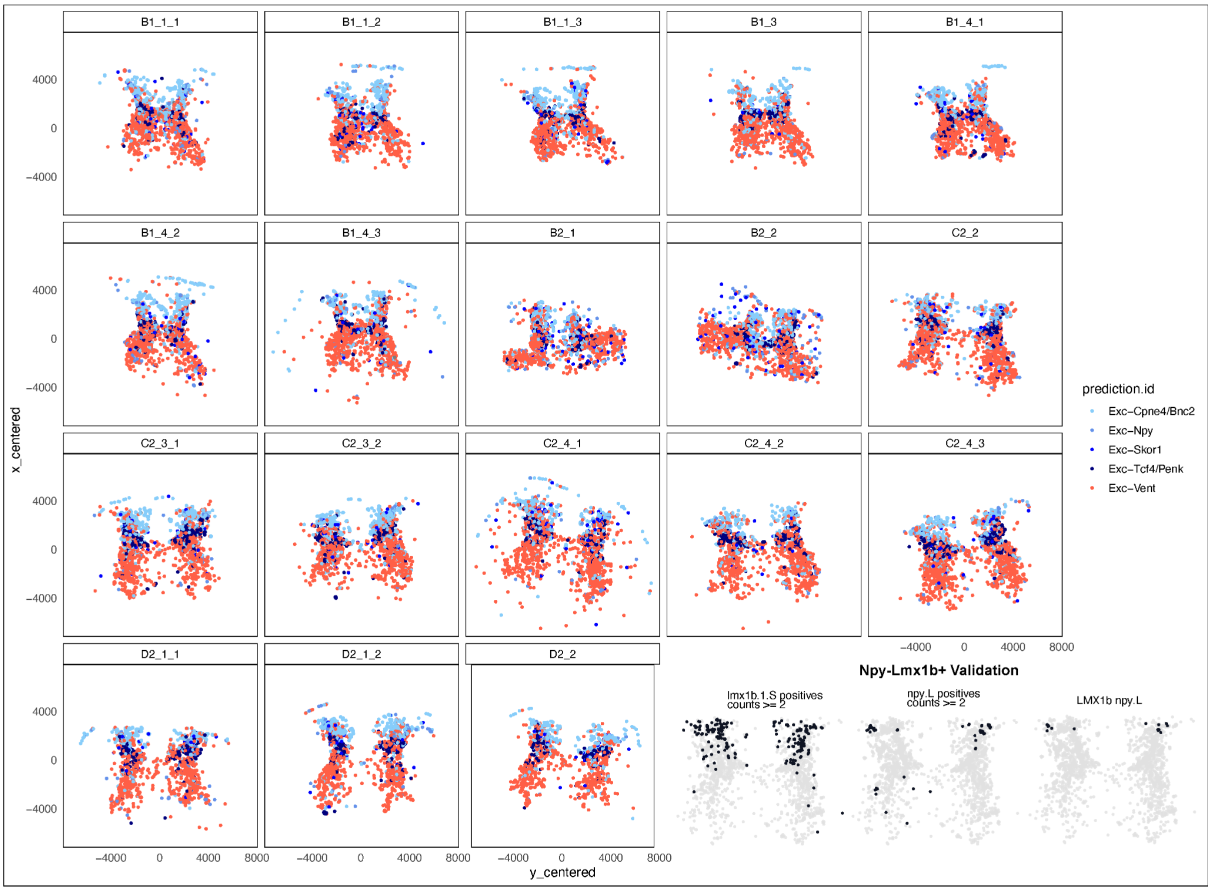
B1_1_1
B1_1_2
B1_1_3
B1_3
B1_4_1
B1_4_2
B1_4_3
B2_1
B2_2
C2_2
C2_3_1
C2_3_2
C2_4_1
C2_4_2
C2_4_3
D2_1_1
D2_1_2
D2_2
x_centered
y_centered
4000
0
−4000
8000
prediction.id
Exc−Cpne4/Bnc2
Exc−Npy
Exc−Skor1
Exc−Tcf4/Penk
Exc−Vent
Npy-Lmx1b+ Validation
lmx1b.1.S positives counts >= 2
npy.L positives counts >= 2
LMX1b npy.L

### Figure S13

**Fig. S13. Representative sections displaying excitatory cell types in the adult frog spatial dataset.** Neuronal identities in the spatial dataset were assigned by label transfer from the single-nucleus reference dataset.

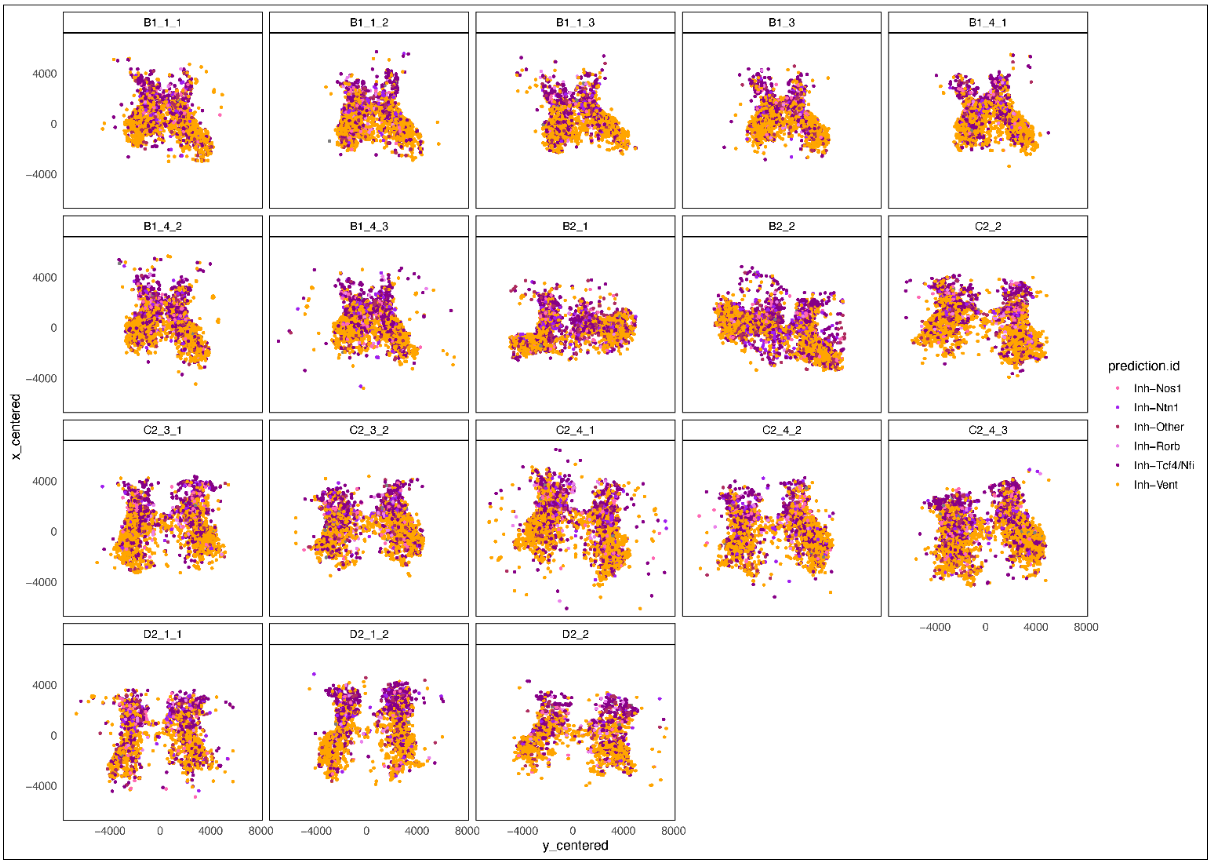
B1_1_1
B1_1_2
B1_1_3
B1_3
B1_4_1
B1_4_2
B1_4_3
B2_1
B2_2
C2_2
C2_3_1
C2_3_2
C2_4_1
C2_4_2
C2_4_3
D2_1_1
D2_1_2
D2_2
x_centered
y_centered
4000
0
−4000
8000
prediction.id
Inh−Nos1
Inh−Ntn1
Inh−Other
Inh−Rorb
Inh−Tcf4/Nfi
Inh−Vent

**Figure S14**

**Fig. S14. Representative sections displaying inhibitory cell types in the adult frog spatial dataset.** Neuronal identities in the spatial dataset were assigned by label transfer from the single-nucleus reference dataset.

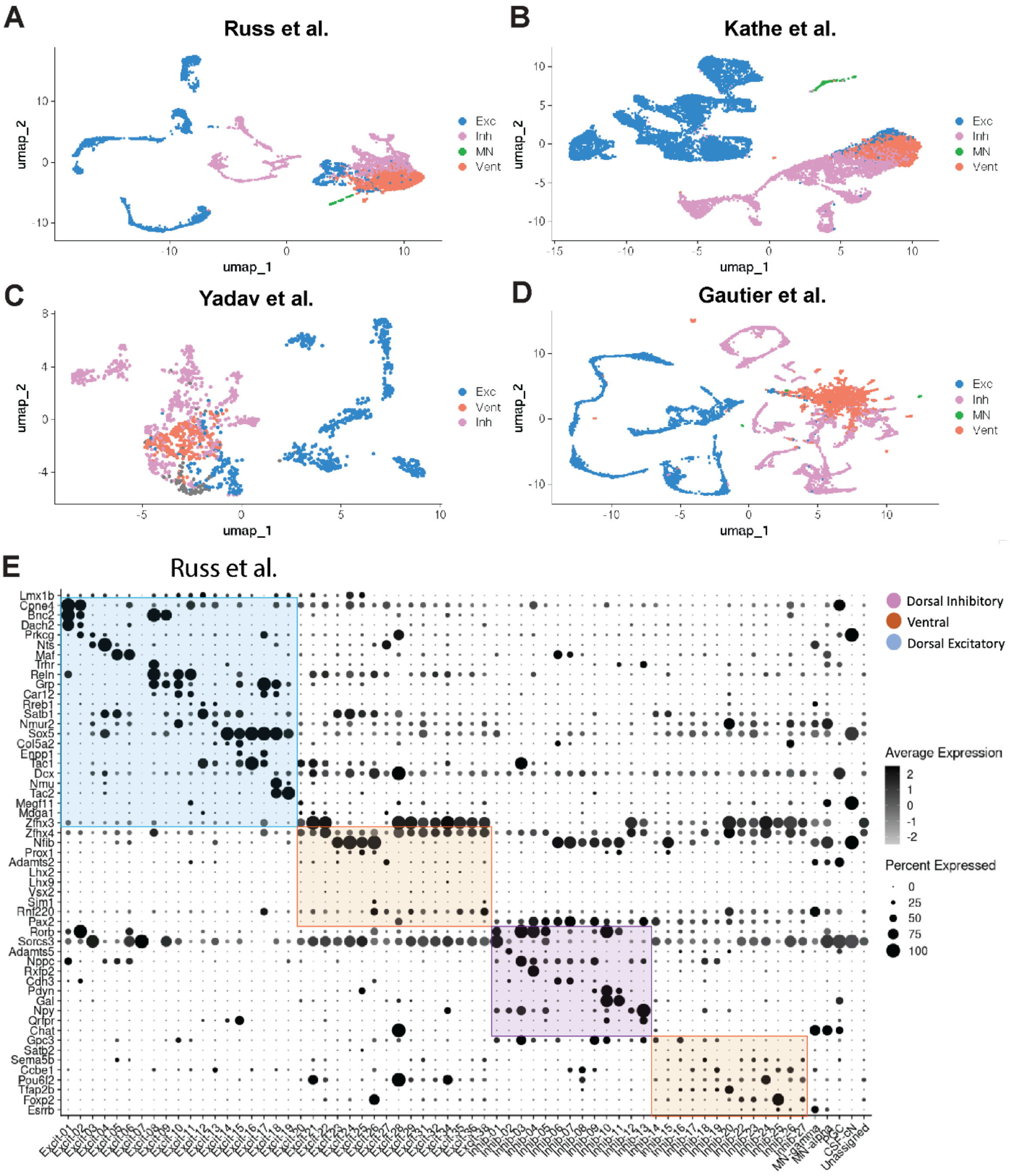
A
Russ et al.
B
Kathe et al.
C
Yadav et al.
D
Gautier et al.
umap_1
umap_2
Exc
Inh
MN
Vent
E
Russ et al.
Dorsal Inhibitory
Ventral
Dorsal Excitatory
Average Expression
Percent Expressed

### Figure S15

**Fig. S15. Analysis of published adult mouse and human single nuclei datasets.**
**A-D.** UMAP representation of single-nuclei datasets for mouse (**A,B**) and human (**C,D**) adult spinal cord colored by coarse neural type group.
**E.** Dot plot showing normalized expression of neural type marker genes in Russ et al.

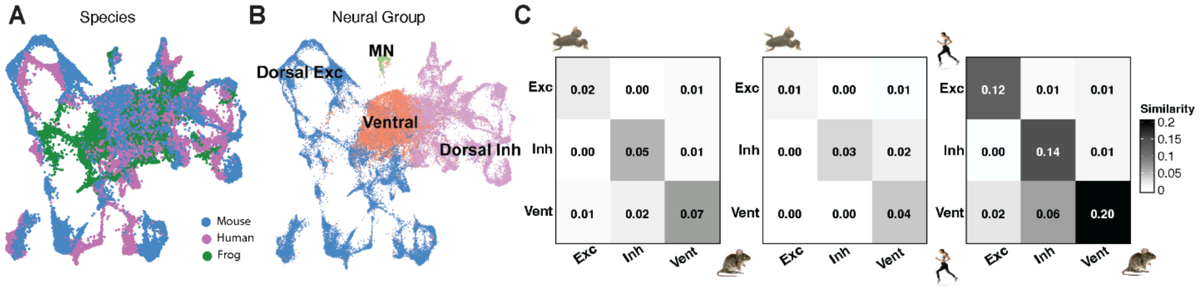
A
Species
Mouse
Human
Frog
B
Neural Group
MN
Dorsal Exc
Ventral
Dorsal Inh
C
Exc
Inh
Vent
0.02
0.00
0.01
0.00
0.05
0.01
0.01
0.02
0.07
Exc
Inh
Vent
0.01
0.00
0.01
0.00
0.03
0.02
0.00
0.00
0.04
Exc
Inh
Vent
0.12
0.01
0.01
0.00
0.14
0.01
0.02
0.06
0.20
Similarity
0.2
0.15
0.1
0.05
0

### Figure S16

**Fig. S16. RPCA integration of the adult neuronal data from frog, mouse and human.**
**A-B.** UMAP-representation of frog, mouse and human RPCA-integrated neural adult data labeled by species (**A**) and coarse neural type group (**B**).
**C.** Heatmaps depicting pairwise similarity indices among coarse neural type groups within the integrated frog–mouse–human dataset, with each panel representing a comparison between two species.

*Cross-species similarity of neural groups in development*

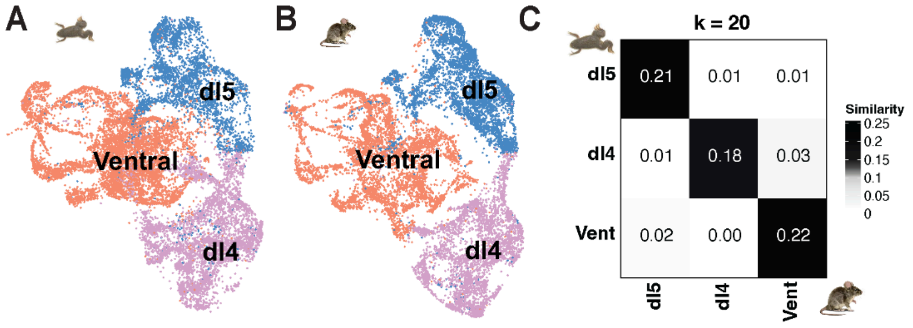


### Figure S17

**Fig. S17. Cross-species integration and similarity analysis of developing NF stage 54 frog and embryonic E9.5-13.5 mouse data, analogous to adult frog and mouse comparisons in Fig. 4F.**

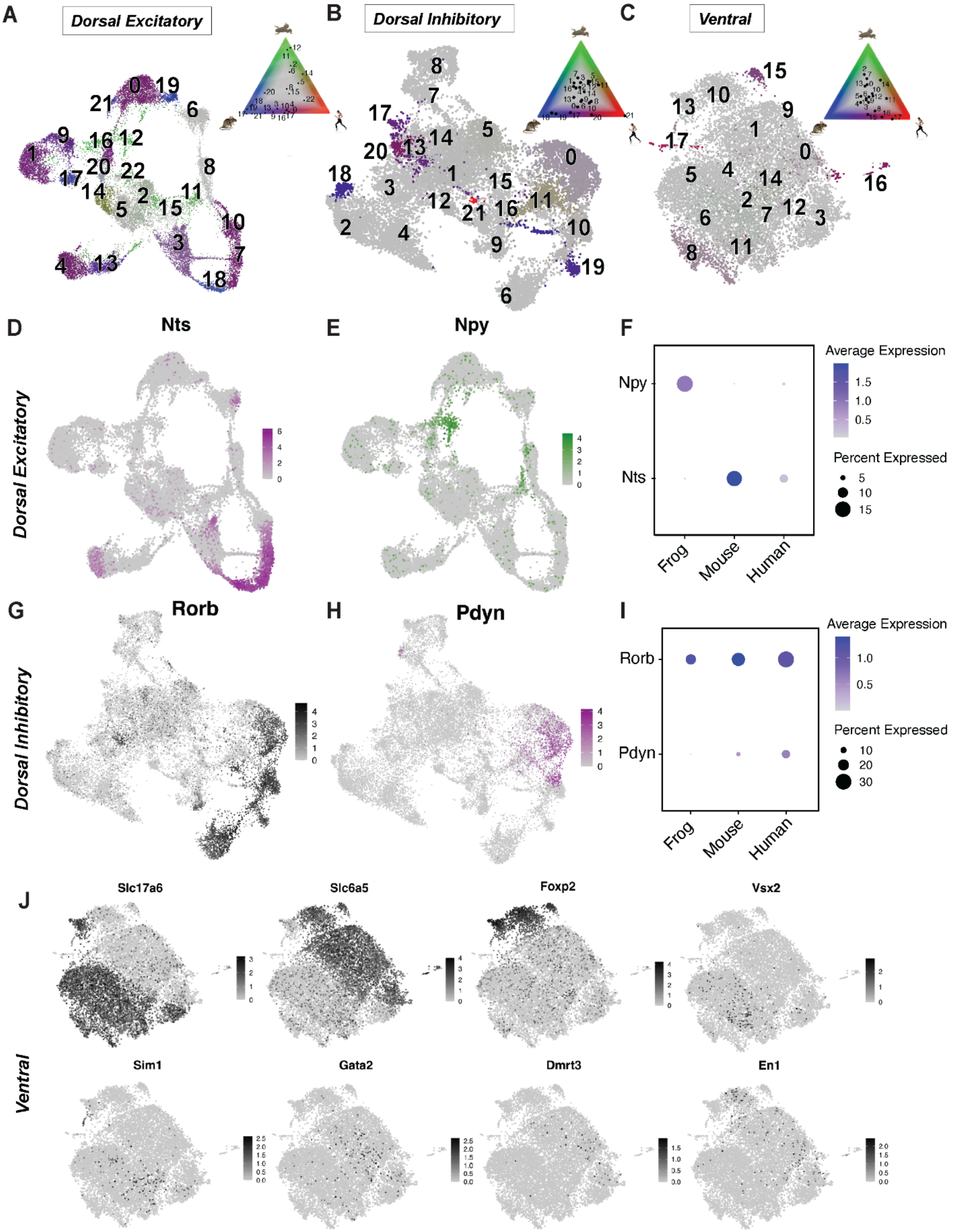
A
Dorsal Excitatory
B
Dorsal Inhibitory
C
Ventral
D
Nts
E
Npy
F
Average Expression
Percent Expressed
Npy
Nts
Frog
Mouse
Human
G
Rorb
H
Pdyn
I
Average Expression
Percent Expressed
Rorb
Pdyn
Frog
Mouse
Human
Dorsal Excitatory
Dorsal Inhibitory
J
Slc17a6
Slc6a5
Foxp2
Vsx2
Sim1
Gata2
Dmrt3
En1
Ventral

**Figure S18**

**Fig. S18. Three-species integration expanded.**

**A-C.** Integrated UMAP representations of frog, mouse, and human neurons for dorsal excitatory (**A**), dorsal inhibitory (**B**), and ventral (**C**) neurons, color-coded by species-mixing values. Inset depicts mixing values across populations with red color representing human, blue - mouse, and green - frog.

**D-E.** Feature plots demonstrating the normalized expression of neural type markers such as amphibian-specific *Npy* and mammalian specific *Nts* within dorsal excitatory integrated neurons.

**F**. Dotplot showing the normalized expression of *Npy* and *Nts* markers within integrated data split by frog, mouse and human.

**G-H.** Feature plots demonstrating the normalized expression of neural type markers such as conserved *Rorb* and mammalian-specific *Pdyn* within dorsal inhibitory integrated neurons.

**I**. Dotplot showing the normalized expression of *Rorb* and *Pdyn* markers within integrated data split by frog, mouse and human.

**J.** Feature plots demonstrating the normalized expression of neural type markers across ventral integrated neurons.

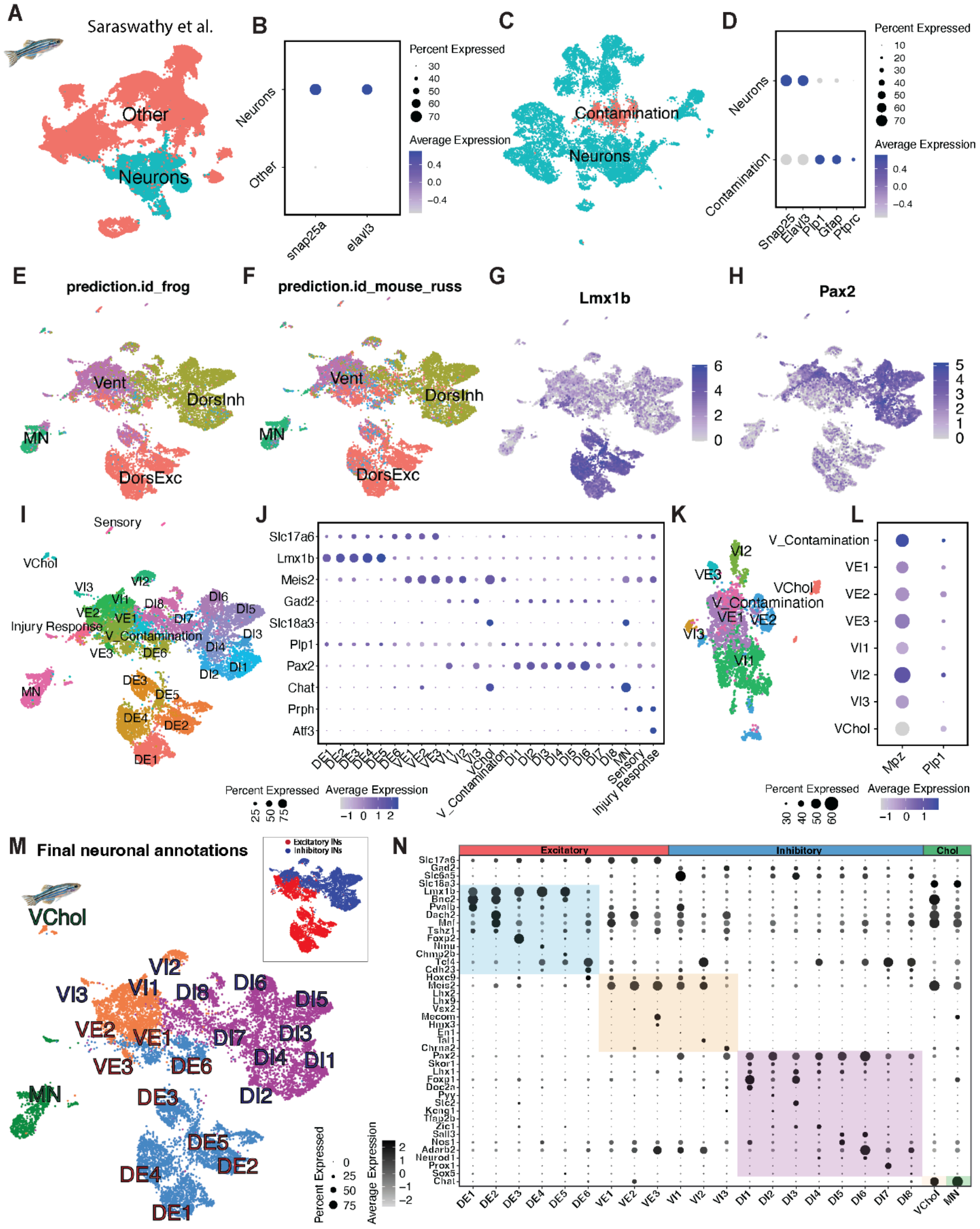

A
Saraswathy et al.
Other
Neurons
B
Percent Expressed
Average Expression
C
Contamination
Neurons
D
E
prediction.id_frog
F
prediction.id_mouse_russ
G
Lmx1b
H
Pax2
I
J
K
L
M
Final neuronal annotations
N
Excitatory
Inhibitory
Chol


### Figure S19

**Fig. S19. Analysis of adult zebrafish spinal cord neural data from *Saraswathy et al. 2024.***

**A.** UMAP representation displaying zebrafish single-nuclei data colored by identity, with neurons in blue and other cellular identities in orange.

**B.** Dotplot displaying neuronal marker genes Snap25a and Elavl3 enriched in neuronal clusters and other cellular identities as represented in **A**.

**C.** UMAP representation displaying subsetted neuronal zebrafish single-nuclei data colored by identity, with neurons in blue and contamination in orange.

**D.** Dot plot displaying neuronal marker genes Snap25a and Elavl3 and other genes (Plp1, Gfap, Ptprc) in neuronal clusters and contamination, showing mutually exclusive expression, as represented in **C**.

**E-H.** UMAPs display filtered neuronal zebrafish single-nuclei data colored by label-transfer neuronal identity (dorsal inhibitory, DorsInh, yellow; dorsal excitatory, DorsExc, red; ventral, Vent, pink; or motor neurons, MNs, green) when the reference is frog (**E**) or mouse (**F**); or by the dorsal excitatory marker Lmx1b (**G**) and inhibitory marker Pax2 (**H**).

**I.** Same data as in **E-H** colored and numbered according to unbiased clusters with contaminating and injury response clusters indicated

**J–L.** Dot plot showing expression across the unbiased clusters from **I,** before removal of contaminating glial cells (**K-L**).

**M.** UMAP representation of the filtered final neuronal dataset, color-coded by unbiased clusters. Inset, same neurons color-coded by excitatory (red) or inhibitory (blue) classification.

**N.** Dot plot showing marker gene expression across unbiased clusters in the filtered neuronal dataset.

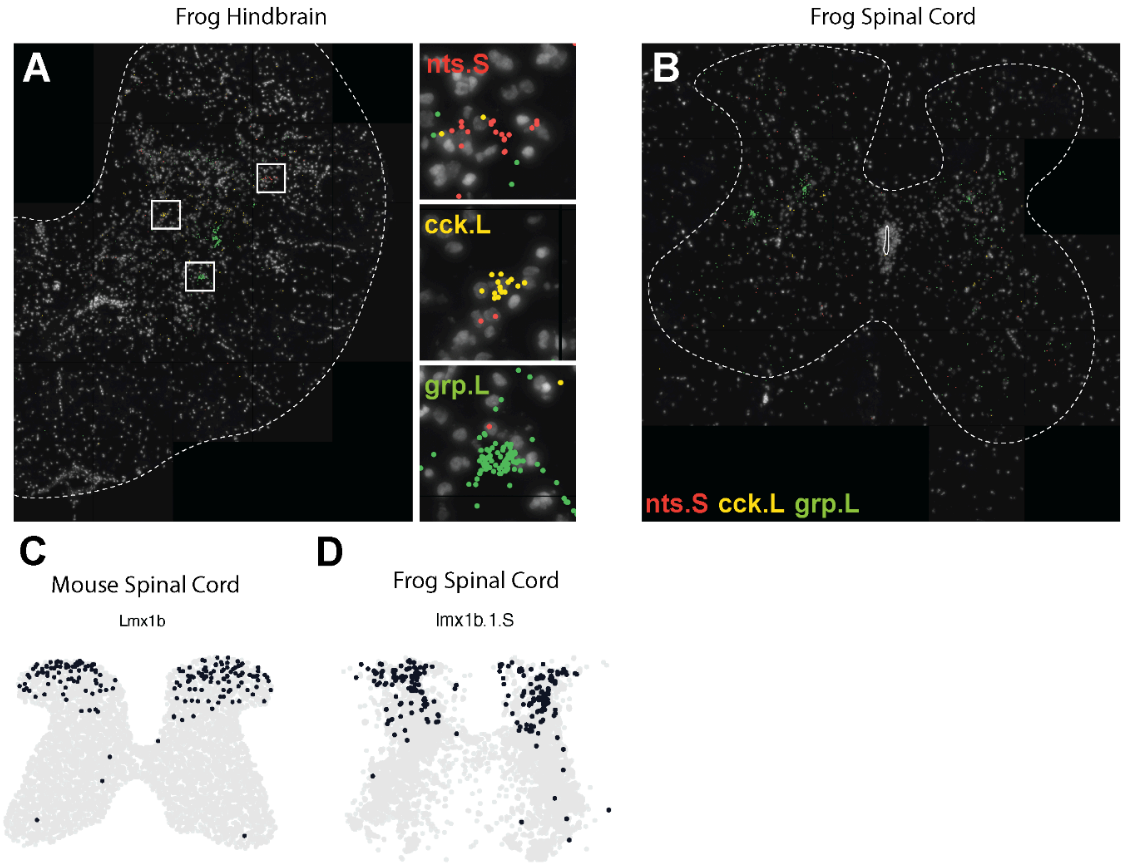
Frog Hindbrain
A
nts.S
cck.L
grp.L
Frog Spinal Cord
B
nts.S cck.L grp.L
C
Mouse Spinal Cord
Lmx1b
D
Frog Spinal Cord
lmx1b.1.S

**Figure S20**

**Fig. S20. Extended spatial transcriptomic validation analyses of mammalian-specific dorsal excitatory subtypes in adult mouse and frog spinal cord.**

**A-B.** Neuropeptides Nts (red), Cck (yellow) and Grp (green) are expressed in the adult hindbrain (**A**), validating the efficacy of our *in situ* RNA expression probe sets (Resolve). In the spinal cord, Grp (green) is expressed in the ventral spinal cord but Nts (red) and Cck (yellow) are largely absent from the dorsal horn (**B**).

**C-D**. Dorsal excitatory Lmx1b-positive cells in the adult mouse (**C**) and frog (**D**) spinal cord.
